# Senolytic Treatment for Low Back Pain

**DOI:** 10.1101/2024.01.15.575738

**Authors:** Matthew Mannarino, Hosni Cherif, Saber Ghazizadeh, Oliver Wu Martinez, Kai Sheng, Elsa Cousineau, Seunghwan Lee, Magali Millecamps, Chan Gao, Jean A. Ouellet, Laura Stone, Lisbet Haglund

## Abstract

Senescent cells (SnCs) accumulate due to aging and external cellular stress throughout the body. They adopt a senescence-associated secretory phenotype (SASP) and release inflammatory, and degenerative factors that actively contribute to age-related diseases such as low back pain (LBP). The senolytics, o-Vanillin and RG-7112, remove senescent human intervertebral (IVD) cells and reduce SASP release, but it is not known if they can treat LBP. *sparc^-/-^* mice, with LBP, were treated orally with o-Vanillin and RG-7112 as single or combination treatments. Treatment reduced LBP and SASP factor release and removed SnCs from the IVD and spinal cord. Treatment also lowered degeneration score in the IVDs, improved vertebral bone quality, and reduced the expression of pain markers in the spinal cord. The result indicates that RG-7112 and o-Vanillin with the combination treatment providing the strongest effect are potential disease-modifying drugs for LBP and other painful disorders where cell senescence is implicated.

**One Sentence Summary:** Senolytics drugs can reduce back pain

## INTRODUCTION

Senescent cells (SnCs) accumulate in ageing and degenerating tissues and are thought to directly contribute to disorders including heart disease, cancer(*1–3*), and osteoarthritis(*4–6*). Age-related accumulation is due to successive shortening of telomere length during replicative cycles(*7, 8*) while stress-induced senescence can be induced prematurely by stressors such as DNA damaging agents, oxidative stress, mitochondrial dysfunction, load-induced injury and disruption of epigenetic regulation (*9, 10*). SnCs are resistant to apoptosis and, in addition to changes in their replicative status, release an array of inflammatory cytokines, chemokines, and proteases known collectively as the *senescence-associated secretory phenotype (SASP)*(*1*). The inflammatory environment triggered by SnCs prevents adjacent cells from maintaining tissue homeostasis(*11, 12*), and it induces senescence in a paracrine manner, thus exacerbating tissue deterioration(*13*). All SnCs share these general features, but there are distinct differences in SASP and anti-apoptotic pathways linked to cell type, species, and inducer of senescence(*14, 15*).

Drugs that selectively target and remove SnCs (senolytic) have recently been identified(*16, 17*). There are four main groups described to date targeting specific pro-senescence and anti-apoptotic pathways. 1) Inhibitors of the Bcl-2 family of apoptosis regulatory proteins. 2) Inhibitors of the p53/MDM2 complex, that alleviate resistance to apoptosis. 3) HSP-90 and PI3K/Akt inhibitors, releasing pro-apoptotic transcription factors. 4) Natural flavonoids with a less clear mode of action(*2, 4, 16, 18, 19*). Targeting indirectly related pro-senescence and anti-apoptotic pathways has shown promising results with increased selectivity for SnCs in the absence of toxicity for normal proliferating or quiescent cells, as exemplified by the combination of dasatinib (D) and quercetin (Q) for age-dependent IVD degeneration and age-related pathologies in mice(*20–24*).

Low back pain (LBP) is often related to IVD degeneration and is the number one global cause of years lived with disability(*25–28*). The personal costs of reduced quality of life and the economic cost to healthcare systems are enormous(*29*), exceeding $100 billion per year in the US alone(*30*).SnCs accumulate in degenerating IVDs and are proposed to directly contribute to disease progression and back pain(*31–33*). The Senolytic drug RG-7112 is a p53/MDM2 complex inhibitor. o-Vanillin is a natural senolytic and senomorphic substance that has been shown to reduce senescence burden and SASP factor release and to improve tissue homeostasis in human IVDs(*34–36*)indicating that they could potentially reduce pain.

The primary manifestation of IVD morbidity in humans is LBP, which cannot be evaluated in cell and tissue culture experiments. Here, we turned to the *sparc^-/-^* mouse model to determine if RG-7112 and o-Vanillin can attenuate behavioral indices of pain and IVD degeneration without causing adverse effects. SPARC (Secreted Protein Acidic and Rich in Cysteine) is one of the most downregulated genes in IVDs of patients with LBP, and mice lacking the *sparc* gene develop age-related IVD degeneration and LBP(*37–39*). The primary objective of this study was to determine if o-Vanillin and RG-7112 remove SnCs, reduce inflammatory mediators, and relieve pain in middle-aged *sparc^-/-^*mice with established IVD degeneration and back pain. The secondary objective was to evaluate if a combination of the drugs provides a better effect than either drug alone.

## RESULTS

### Validation of the *sparc^-/-^* mouse model to study back pain and cell senescence

SnCs are suggested to contribute to IVD-related LBP in human patients, and removing SnCs in human IVD tissue and cell cultures decreases the expression of inflammatory and pain-mediating SASP factors. Human patients with LBP have reduced gene expression, epigenetic modifications, and lower SPARC protein content in degenerating IVDs(*40, 41*). Similarly, animals with the *sparc gene* depleted develop age-related LBP, and like in human patients, some areas of the spine are more affected than others (**Fig.1(a-b)**). We found that *sparc^-/-^* mice, like humans, accumulate SnCs in their IVDs with age and degeneration. The NP region had a 3-fold and the AF region about 6-fold increase in *p16^Ink4a^* positive SnCs when comparing age-matched *sparc^-/-^* and wildtype IVDs (**Fig.1(c-d)**). An increase in the level of SnCs and the degree of degeneration in mice IVDs (L3-S1) were strongly correlated in both NP (R^2^=0.9) and AF (R^2^=0.95) regions (**Fig.1e**).

**Fig.1.**
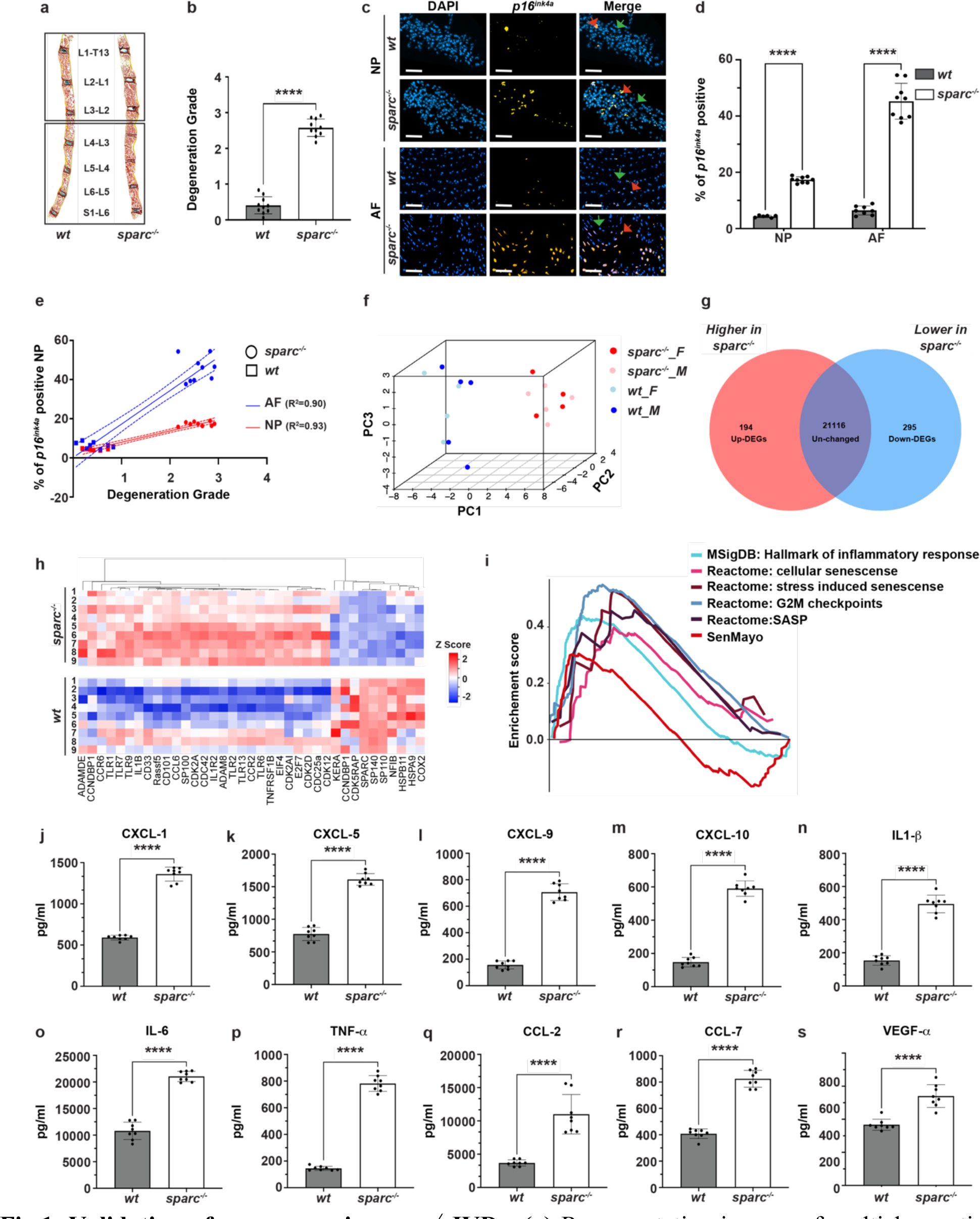
Validation of senescence in *sparc^-/-^* IVDs. **(a)** Representative images of multichromatic FAST stained 16 mm sections of 9-month-old *sparc^-/-^* and wild-type spines (T13-S1). **(b)** Quantification of the histological degeneration grade in IVDs of the lower lumbar spine where degeneration is frequently found (L3-S1) is shown in **(a)**. n = 10 animals per group (5 males and 5 females). Each value represents the average grading score of 3-4 IVDs per animal. **(c)** Photomicrographs showing *p16^Ink4a^* positive NP and AF cells with DAPI counterstain of *sparc^-/-^* and wild-type IVDs (L3-S1). The red arrows point to *p16^Ink4a^* positive cells, and the green arrows indicate non-senescent cells. **(d)** Quantifying the percentage of *p16^Ink4a^*-positive cells shows increased numbers in the NP and AF of *sparc^-/-^* IVDs compared with wild-type. **(e)** Scatter plot shows a positive correlation between IVD degeneration and the number of *p16^Ink4a^*positive NP (red line) and AF (blue line) cells both in wild-type and *sparc^-/-^* mice. Each value represents the average percentage of *p16^Ink4a^* positive cells from 3-4 IVDs per animal of 6-8 WT and 9 *sparc^-/-^* animals per group in (**c-d**). **f.** PCA analysis demonstrated a clear difference in transcriptomic signatures of upper lumbar IVDs without clear degenerative changes (T13-L3) of *sparc^-/-^* and WT mice. **g.** Venn diagram depicting significant differentially up and downregulated genes in *sparc^-/-^*compared to WT mice. **h.** Heatmap of 37 significantly differentially expressed senescence and SASP-associated genes in *sparc^-/-^* and WT discs. Data shown are relative to the calculated Z scores across the samples and ranked by significance adjusted to p<0.05. Red represents relatively high levels of expression; blue represents relatively low levels of expression. Each column represents one animal, and each row represents the expression of a single gene. **i.** Multiplex GSEA of the SenMayo, Reactome (cellular senescence, SASP, Stress-induced senescence, G2M checkpoints) and inflammatory response (MSigDB) gene sets demonstrate a significant accumulation of senescence-associated genes in *sparc^-/-^* IVDs compared with WT. Nominal p-value, calculated as a two-sided t-test, with no adjustment since only one gene set was tested. n = 9 animals per strain (4 males and 5 females) and 3 pooled IVDs (T13-L3) per animal for a total of 27 IVDs per strain in (**e-i**). IVDs from the lower lumbar level where degeneration is frequently were isolated and the release of 15 SASP factors was evaluated *ex vivo. Sparc^-/-^* IVDs showed a higher release than WT of 10 factors (CXCL-1, -5, -9, -10, IL1-β, IL-6, TNF-α, CCL2, CCL7, VEGF-α). ****p < 0.0001 indicates a significant difference between *wt* and *sparc^-/-^*. n = 4 discs per animal with 8 animals (4 males and 4 females) in each group (Total of 32 IVDs). Values are presented as Mean ± SD and significance was measured by two-tailed unpaired t-tests in (**b** and **j-s**) and repeated measures of one-way ANOVA in (**d**).

We next performed bulk RNA sequencing (RNA seq) to evaluate whether senescence-associated genes were enriched in IVDs of *sparc^-/-^* mice. We used the IVDs from the upper lumbar spine (T13-L3) from wild-type and *sparc^-/-^* mice to minimize the number of animals. Degenerative changes are less frequent in this area compared to the lumbar levels (L3-S1) used for histological analysis. Spearman correlation was computed to examine the variance and the relationship of global gene expression across the samples. Principal component analysis (PCA) shows a clear separation between wild-type and *sparc^-/-^* (**Fig.1f**). We identified 489 DEGS, 194 significantly upregulated, and 295 significantly downregulated in *sparc^-/-^* mice (**Fig.1g**). A significant enrichment was detected in genes regulating senescence (*cdk2a, cdk2ai*(*p16^Ink4a^*)*, E2F7, EIF4, CDC25a, CDC42, cdk2d, cdk12 and TNFRSF1b)* and SASP factors (*IL1-b, IL1R2*, *CCL6, CCR6,* ADAM8, *TLR-1,-2, -7,* and *TLR9)* (**Fig.1h**). Gene set enrichment analysis (GSEA) comparing our data to those presented by the Human Molecular Signatures Database (MSigDB), Reactome and SenMayo, revealed an upregulation of pathways involved in senescence (NES=1.52 and 1.02), G2M cell cycle checkpoints (NES=2.62), and SASP factors release (NES=1.92) in *sparc^-/-^* animals. (**Fig.1i**). To evaluate the influence of senescence, a multiplex assay was used to measure the release of fifteen SASP factors from lumbar discs (L4-S1) where degeneration in *sparc^-/-^* animals is prominent. Ten of the fifteen SASP factors (CXCL-1,-5, -9, -10, IL1-b, IL-6, TNF-a, CCL-2, -7, and VEGF-a) assessed showed significantly higher release in *sparc^-/-^* compared to wild-type IVDs. Three factors (IL-2, -10, and MCSF) did not show a difference, and two factors (INF-γ, and RANKL) were undetected (**Fig.1(j-s)** and **Table S1**). Together this information provides the rationale for evaluating senotherapeutics as a potential treatment for LBP in this model.

The senolytic drugs o-Vanillin and RG-7112, remove senescent human IVD cells and reduce SASP factor release from cells and intact IVDs(*34, 35, 42, 43*). The phenotype of SnCs is heterogeneous and can differ between species, therefore before venturing *in vivo* and to evaluate if o-Vanillin and RG-7112 can also target SnCs mouse IVD cells and reduce SASP factors, isolated IVDs were treated in culture *ex vivo* with single or combined treatments. The selected drug concentrations were based on previous studies with human IVD cells, and intact human IVDs(*34, 35, 42, 43*). The ten factors that were elevated in *sparc^-/-^* were all reduced, with nine out of ten significantly reduced, following treatment with either o-Vanillin or RG-7112. The ten factors were further reduced with a combination of the drugs at the highest concentration (**Fig.S1(a-m) and Table S1**). This data demonstrates that o-Vanillin and RG-7112 are efficacious in mouse IVDs and can potentially affect IVD degeneration and back pain if they reach their target. Since it is not possible to evaluate if treatment can reduce perceived pain in *ex vivo* studies, our next step was to evaluate if treatment can reduce perceived pain.

### Targeting cell senescence with senolytic drugs reduced low back pain in *sparc^-/-^* mice

*Sparc^-/-^* mice start showing signs of LBP at 4 months of age and have well-established pain by 7 months(*37–39*). We compared male and female *sparc^-/-^* and wild-type animals over 8 weeks (7-9 months of age) as a baseline for pain behaviour. Grip strength and tail-suspension were used to determine axial pain, while von Frey and acetone-evoked behaviour were used to evaluate radiating pain(*37–39, 44, 45*). The experimental design and treatments are depicted in **Fig.2a**. As expected, *sparc^-/-^* animals showed well-established behavioural signs of LBP that progressively worsened over the 8-week period, whereas wild-type animals showed a stable pain phenotype (**Fig.2** **(b-i)**). *Sparc^-/-^* animals were treated weekly, by oral gavage to establish if oral treatment with o-Vanillin and RG-7112 reduced LBP. The concentrations were extrapolated from the *ex vivo* studies by converting the highest optimal concentration in culture (µM) to the highest dose *in vivo* (mg/kg)(*46–48*) as outlined in **Table 1**. Both drugs improved the pain phenotype and significantly improved axial discomfort (grip strength and tail suspension), cold sensitivity (acetone-evoked behaviour) and radiating pain (Von Frey) after 4 weeks of treatment, with all assessments significantly improving after 8 weeks of treatment (**Fig.2(b-i)**). To determine if the drugs provide synergistic or additive effects, they were combined at 100+100% (HO+HR), 100+50% (HO+LR), 50+100% (LO+HR), or 50+50% (LO+LR) of the single drug concentrations. After 8 weeks of treatment, combining the drugs while keeping at least one at 100% further significantly improved the effect. The attenuation of pain behaviour was lost in all tests except tail suspension when lowering the dose of both by 50% (**Fig.2(b-i)**). Male and female animals are shown together; no significant sex differences were found. We found no difference in the weight, mortality, body weight, or distance travelled between the groups (**Fig.S2(a-d)**). The data demonstrates that both drugs can reduce behavioral indices of back pain and that combining them within the higher concentration range further reduces pain.

**Fig.2.**
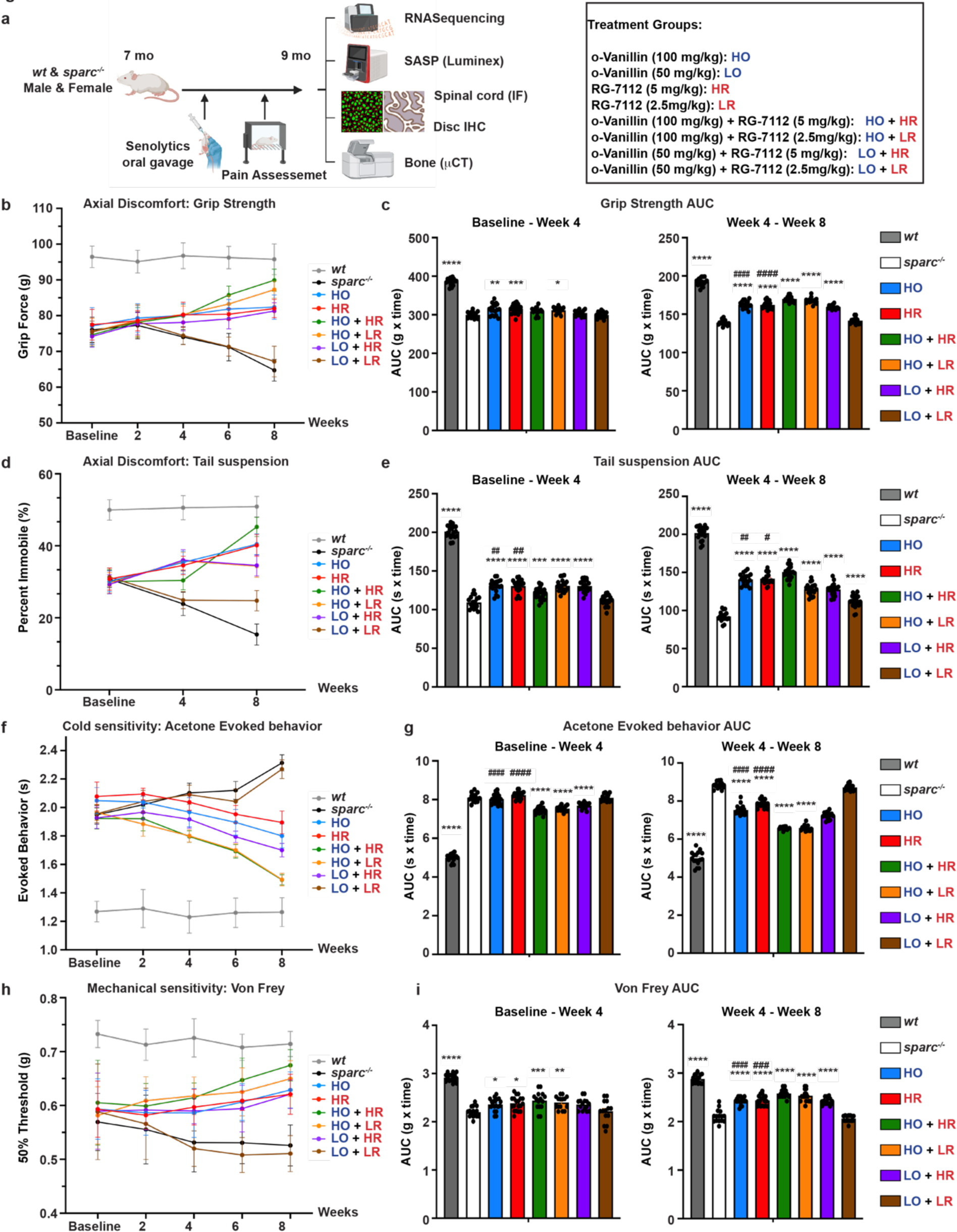
Targeting SCs with senolytic drugs improves axial discomfort, cold, and mechanical sensitivity in *sparc^-/-^* mice. **a.** Schematic of the experimental setup. Animals received vehicle or senolytic treatment weekly through oral gavage (as described in the treatment groups box). Pain assessment tests were performed every 2 weeks (grip strength, acetone-evoked behaviour, von Frey) and every 4 weeks for tail suspension. Axial pain was assessed by **b.** grip strength and **d.** tail suspension tests. Radiating pain was assessed by **f.** acetone-evoked behaviour (cold sensitivity) and **h.** von Frey (mechanical sensitivity) tests . The area under the curve (AUC) between baseline to week 4 and week 4 to week 8 was calculated using the trapezoid method ((T2 − T1) × ((B1 + B2)/2), where T is time and B is the behavioural score for **c.** grip strength, **e**. tail suspension, **g**. acetone-evoked behaviour and **i**. von Frey assay. n = 14–20 animals per group (7-10 males and 7-10 females). Data is presented as mean ± SEM and analyzed by one-way ANOVA followed by Tukey’s post-hoc test (**c**, **e**, **g**, and **i**). * or ^#^ indicates P < 0.05, ** or ^##^ indicates P < 0.01, *** or ^###^ indicates P < 0.001, **** or ^####^ indicates P < 0.0001. * indicates a significant difference compared with *sparc^-/-^* and ^#^ indicates a significant difference compared with single drug treatment.

**Table. 1.** Concentration and annotation for each of the treatment groups.

| Senotherapeutics | Annotation | Concentration (mg/kg) |
| --- | --- | --- |
| o-Vanillin | HO | 100 mg/kg |
| o-Vanillin | LO | 50 mg/kg |
| RG-7112 | HR | 5 mg/kg |
| RG-7112 | LR | 2.5mg/kg |
| o-Vanillin + RG-7112 | HO + HR | 100 mg/kg + 5 mg/kg |
| o-Vanillin + RG-7112 | HO + LR | 100 mg/kg + 2.5 mg/kg |
| o-Vanillin + RG-7112 | LO + HR | 50 mg/kg + 5 mg/kg |
| o-Vanillin + RG-7112 | LO + LR | 50 mg/kg + 2.5 mg/kg |

### Senotherapeutic treatment reduced SASP factor release from the IVDs of *sparc^-/-^* mice

As shown above, IVDs from *sparc^-/-^* animals release measurable and elevated levels of SASP factors that could contribute to LBP (**Fig.1(j-s)**). To determine if the oral treatment reduced SASP factor release *in vivo*, a set of lumbar IVDs was harvested at the termination of the 8-week treatment and the release of the same fifteen SASP factors was measured. Both drugs significantly reduced the release of 10 factors (CXCL-1, -5, -9, -10, CCL-2,-7, IL-1β, -6, TNF-α, and VEGF-α). The combination treatment provided an additive effect and further significantly reduced the release of the same 10 factors. Although significantly lower than untreated, reducing either drug by 50% of the single drug concentrations did not provide any added effect, and reducing both to 50% was less effective than the single drugs at 100% (**Fig.3(a-l)**). Two factors, IL-2, and IL-10 were only significantly decreased by o-Vanillin and not RG-7112 while MCSF was significantly decreased by RG-7112 and not o-Vanillin when single drugs were applied. A significant additive effect was observed for IL-10 and MCSF following combination treatment (**Fig.3(h**, **g**, and **m)**). The measured concentrations and significance are shown in **Table S2**. The data demonstrates that the drugs administered orally could reach the IVDs and reduce SASP factor release.

**Fig.3.**
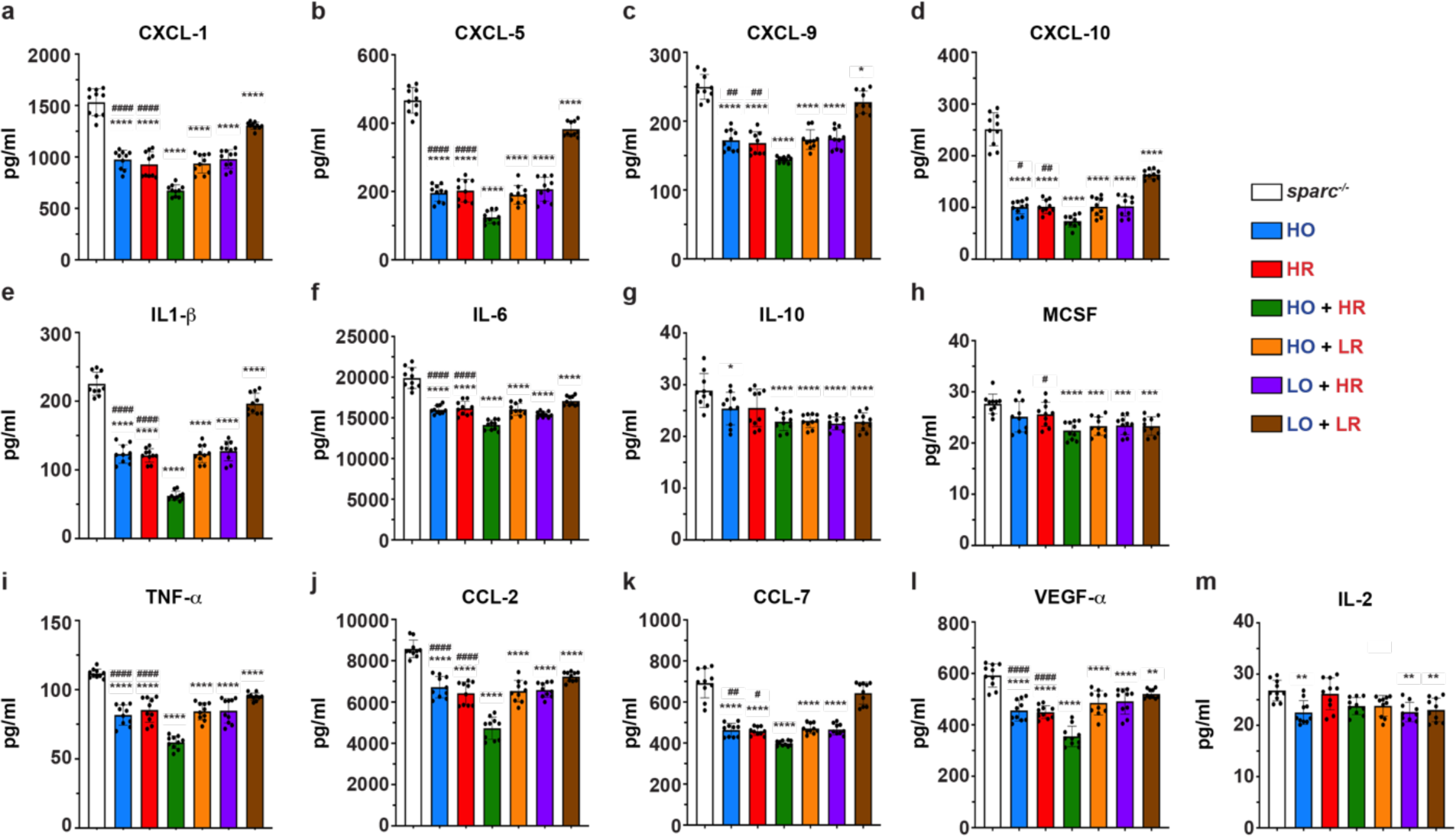
SASP factor release is reduced in treated *sparc^-/-^* mice. IVDs (L3-S1) from *sparc^-/-^* mice treated with senolytics or vehicle were isolated and the release of 15 SASP factors was evaluated. The release of chemokines (**a-d** and **j-k**) cytokines (**e-g** and **i**, **m**), and growth factors (**h** and **l**) was assessed using a multiplex assay. The data is presented as mean ± SD, two-way analysis of variance (ANOVA) and post hoc comparison Tukey’s was used to measure significant differences between the groups. * or ^#^ indicates P < 0.05, ** or ^##^ indicates P < 0.01, *** or ^###^ indicates P < 0.001, **** or ^####^ indicates P < 0.0001. * indicates a significant difference compared with *sparc^-/-^* and ^#^ indicates a significant difference compared with single drug treatment. n = 4 discs per animal for 10 animals (5 males and 5 females) in each group for a total of 40 IVDs per group.

### Senotherapeutics reduced the number senescent IVD cells and improved IVD health in ***sparc^-/-^* mice.**

Next, we evaluated if the number of SnCs *p16^Ink4a^* positive cells was reduced in the IVDs and if treatment could improve IVD health. Single-drug treatment significantly reduced the number of *p16^Ink4a^*-positive cells (∼40%) from both the NP and AF regions (**Fig.4(a-b)**). Combination treatment provided a further significant (25-27%) reduction in both regions when at least one of the drugs was kept at 100%; reducing both by 50 % did not provide an additive effect (**Fig.4(a-b)**). Previous treatment modalities have shown pain reduction in the *sparc^-/-^* model, but no drugs to date have been able to improve histological degeneration score(*37–39, 44, 45, 49–51*). Here we show that both drugs improved histological degeneration score (**Fig.4(c-d)**). Combining the drugs at 100% provided a significant additive effect while lowering one to 50% remained at the level of single drugs and lowering both by 50% failed to improve histological degeneration grade (**Fig.4e**).

**Fig.4.**
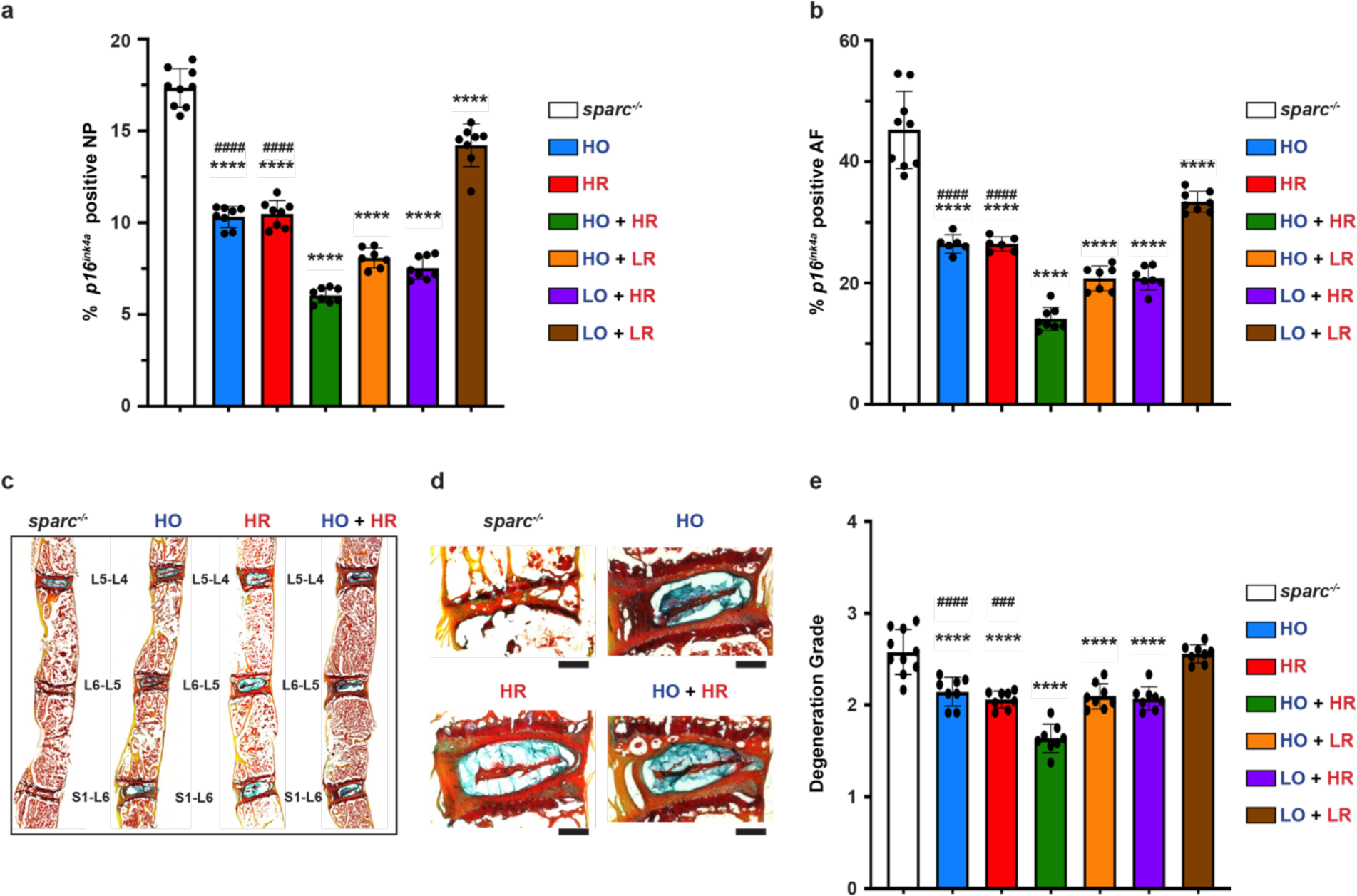
Senolytics remove senescent NP and AF cells and improve disc health. *Sparc^-/-^*IVDS (L3-S1) from animals treated with senolytics or vehicle (n=7-9 animals, 3-5 males and 4-5 females per group) were collected and processed for histological evaluation. Immunofluorescence staining measures the percentage of *p16^Ink4a^*-positive cells, DAPI was used as a counter stain **a.** NP and **b.** AF. **c.** Representative images of lumbar IVDs (L3-S1) from *sparc^-/-^* mice treated with senolytics or vehicle. **d.** FAST staining revealed histological improvement in the ventral region (clear distinction between AF and NP and increase in disc height) of treated discs. **e.** Histological degeneration grade of the IVDs was independently evaluated and the average grade per animal is presented for a total of 8-10 animals (4-5 males and females) and 32-40 IVDs per group. Statistical comparisons were calculated using an ordinary two-way ANOVA, with a Tukey’s post-hoc analysis. Data is presented as mean ± SD. ^###^ indicates P < 0.001, **** or ^####^ indicates P < 0.0001. * indicates a significant difference compared with *sparc^-/-^* and ^#^ indicates a significant difference compared with single drug treatment.

### Senotherapeutics reduced the number senescent cells and pain-signalling in the spinal cord of *sparc^-/-^* mice

SnCs have also been detected in various cell types in the CNS, including neural and glial cells, where their presence could contribute to a pain phenotype(*52–56*). We found a 4.8-fold significantly larger area of *p16^Ink4a^* immunoreactivity in the dorsal horn of *sparc^-/-^* mice compared to wild-type (**Fig.5(a-b)**). Using confocal microscopy, we also confirmed the *p16^Ink4a^* immunoreactivity colocalized with nuclear DAPI staining (**Fig.S3a**). Systemic senotherapeutic treatment could directly affect pain signalling in the spinal cord and contribute to the reduction of pain. Indeed, we found a significant reduction (24-30%) in *p16^Ink4a^*reactivity with single drugs; combining them further significantly reduced the *p16^Ink4a^* -immunoreactivity (30-64%). The effect was lost when both senolytics were reduced by 50% (**Fig.5c**). As previously reported, *sparc^-/-^* mice have increased immunoreactivity for calcitonin gene-related peptide (CGRP), glial fibrillary acidic protein (GFAP) and the activated microglia marker protein (cd11-b) in the dorsal horn of the spinal cord that may contribute to the pain phenotype(*44, 45, 57*) **(Fig.5d**). The immunoreactivity of CGRP was significantly reduced in mice treated with single drugs (52-55%). Combining them further significantly reduced CGRP immunoreactivity (∼60%) with a lower (20%) but significant effect observed even when both were reduced by 50% **(Fig.5e**). Activated microglia and astrocytes in the dorsal horn of the spinal cord are known to contribute to the development of chronic pain(*58*). GFAP immunoreactivity was reduced by single drug treatment (42-44%) with combination treatment keeping the higher o-Vanillin concentration further reducing GFAP immunoreactivity (7-13%) **(Fig.5f**). Lowering o-Vanillin in the combination treatments did not provide an added effect over single drugs. CD11-b immunoreactivity was also significantly reduced with single drugs (40-42%); the effect was further reduced (62%) with the combination treatment at the highest concentration lowering one didn’t provide an additive effect over single drugs and an effect lost when both were given at 50% **(Fig.5g**). These results suggest that senotherapeutics could also be contributing to reducing back pain by decreasing pain-related neuroplasticity and glial cell activity.

**Fig.5.**
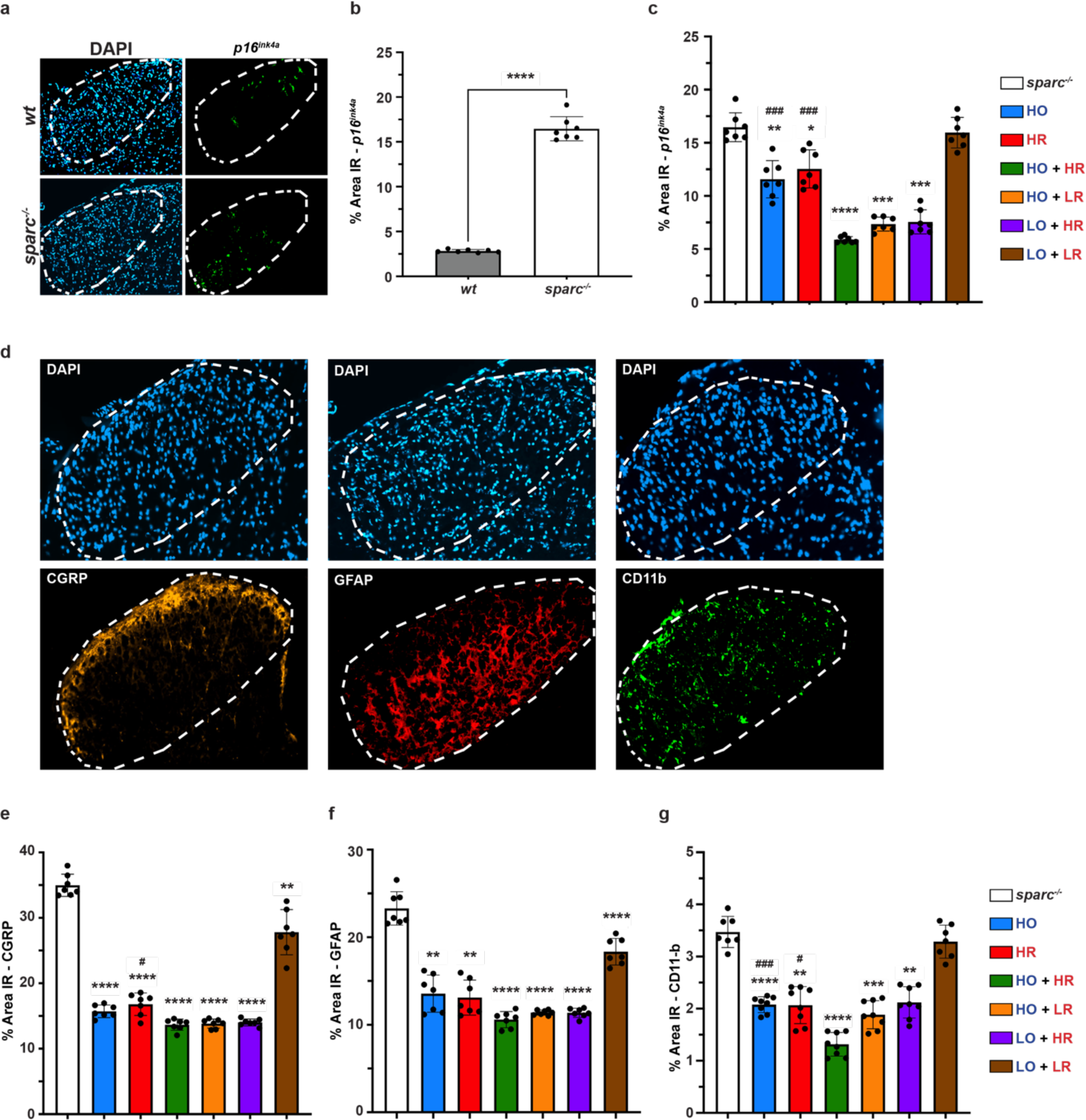
Senolytics reduced SnCs and pain-related neuroplastic changes in the spinal cord of *sparc^-/-^*mice. **a.** Spinal cords from perfused animals treated with senolytic drugs or vehicle were collected and processed for histological evaluation. Representative images showing higher *p16^Ink4a^*-immunoreactivity in the dorsal horn (delineated in dashed line) in *sparc^-/-^* compared to wild-type mice. **b.** Quantification of the increased immunoreactivity (percentage area). **c.** *p16^Ink4a^* immunoreactivity was reduced in treated *sparc^-/-^* animals. The spinal cords were also assessed for pain-related neuroplastic changes. **d.** Representative image of CGRP, GFAP, CD11b immunoreactivity, assessed in the dorsal horn (delineated in a dashed line). The immunoreactivity (percentage area) was quantified for **e.** CGRP, **f.** GFAP, and **g.** CD11b. The % area with immunoreactivity above a set threshold was calculated from the total area of the dorsal horn. n=7-8 animals per group (3-4 males, 4 females). DAPI was used as a counterstain. The average of three separate images was calculated for each animal and used to calculate the mean of the treatment group. Data is presented as mean ± SD and was analyzed by a Two-tailed t-test or an ordinary one-way ANOVA followed by Tukey’s post-hoc test. * or ^#^ indicates P < 0.05, ** indicates P < 0.01, *** or ^###^ indicates P < 0.001, **** indicates P < 0.0001. * indicates a significant difference compared with *sparc^-/-^* and ^#^ indicates a significant difference compared with single drug treatment.

### Senotherapeutics improved bone parameters in *sparc^-/-^* mice

In addition to IVD degeneration and LBP, *sparc^-/-^* mice are known to have osteopenia which is also a potential contributor to LBP. Recent studies have demonstrated that senolytic drugs can improve age-related bone loss. We therefore treated a second cohort of animals to evaluate if oral administration of o-Vanillin and RG-7112 could improve bone quality in *sparc^-/-^* mice. We treated *sparc^-/-^* animals with single and combination treatments, keeping the drug concentrations at higher levels (HO, HR, and HO+HR). Micro-CT analysis was performed, and we evaluated IVD volume and vertebral bone quality. Previous studies have shown a reduced disc height index in *sparc^-/-^*compared to wild-type mice and our analysis demonstrated a significantly lower IVD volume (∼34%) in the *sparc^-/-^* group (**Fig.6(a-b)**). Treatment with the two drugs as single treatments showed a trend towards improvement in disc volume compared to vehicle-treated mice. In contrast, a significant increase (∼27%) in disc volume was found following combination treatment with both drugs at 100% (**Fig.6c**), **Fig. S4(a-b)**, and **Movie S1-5**. 3D reconstructions representing vertebrae of treated and untreated animals are illustrated in **Fig.6d**, **Fig.S4(a-b)**, and **Movie S1-5**. Analysis of bone microarchitecture was performed in a region starting just above the growth plate to the beginning of the pedicels as illustrated in (**Fig.S4b**). The trabecular microarchitecture revealed a significantly lower bone density (56%), trabecular thickness (15%), and trabecular number (50%) in *sparc^-/-^* compared to wildtype mice (**Fig.6(g-i)**). In contrast, trabecular separation was significantly higher (42%) in *sparc^-/-^*compared to wild-type mice (**Fig.S4c**). Single treatment of *sparc^-/-^* animals resulted in a significantly increased bone density, BV/TV, (52% and 69%) that was somewhat stronger (85%) and more significant by the combination treatment (**Fig.6j**). Similarly, each drug alone or combination treatments resulted in a significant increase in trabecular number (34.1%, 48%, and 47%) (**Fig.6k)**. Trabecular thickness was significantly increased only following treatment with RG-7112 and combination (15%, and 23%) (**Fig.6l)**. No significant improvement was observed in trabecular separation for any of the treatment groups (**Fig.S4d**).

**Fig.6.**
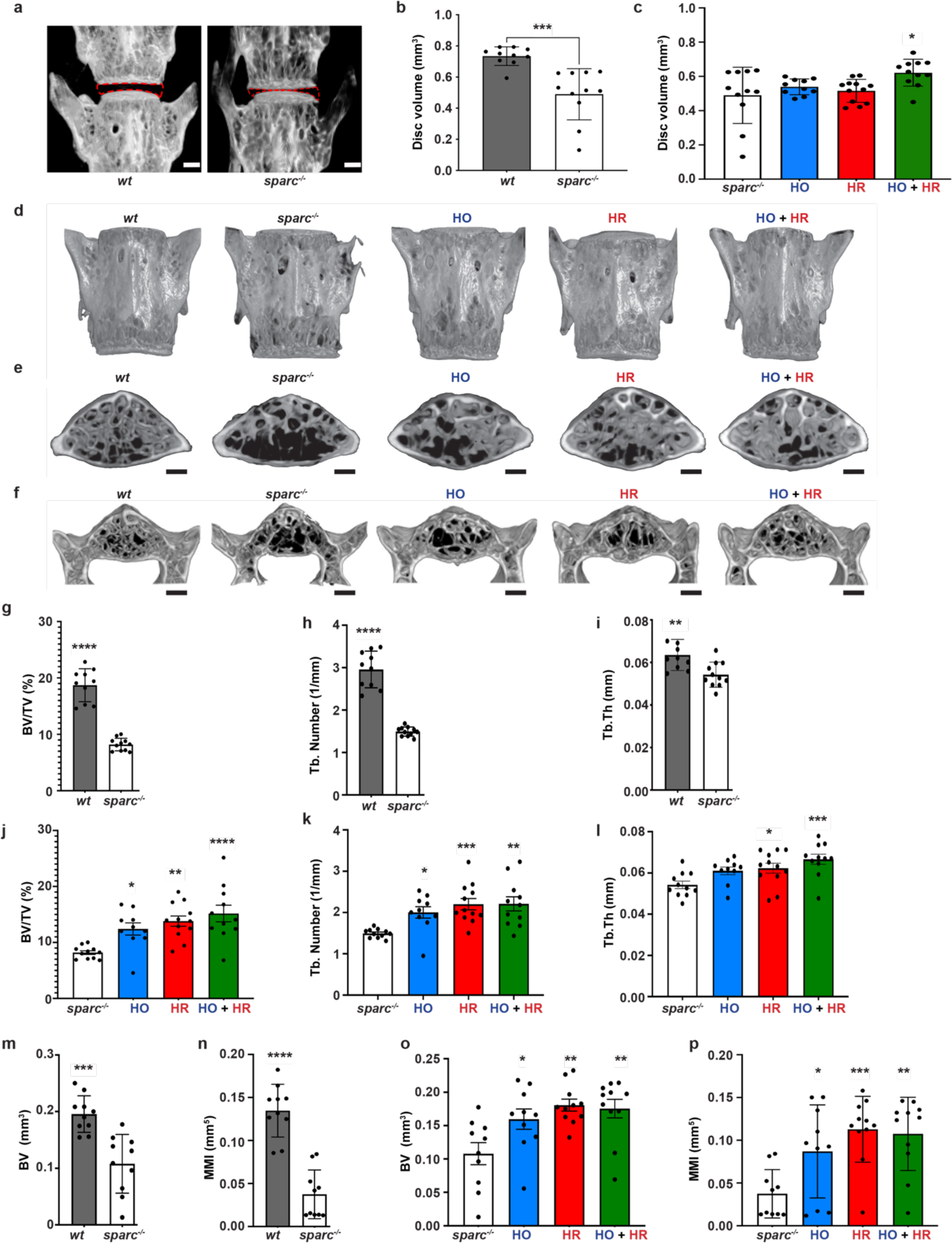
Treatment with senolytics resulted in increased IVD volume and improved bone quality. **a.** Representative images showing the difference in disc volume (delineated in red) between *sparc^-/-^* and wild-type mice. **b.** Quantification of disc volume showed a lower volume in *sparc^-/-^* compared to WT mice. **c.** Disc volume of *sparc^-/-^* IVDs from animals treated with senolytics or vehicle. Representative μCT images showing **(d)** vertebrae, **(e-f)** trabecular and cortical bone of wild-type and *sparc^-/-^* mice (vehicle and treated). Quantification of trabecular **(g-l)** and cortical **(m-p)** bone parameters. Scale bar in **a**, **e**, and **f** = 500 µm. Statistical comparisons were calculated using a Two-tailed t-test or an ordinary one-way ANOVA, with a Dunnett’s post-hoc analysis as appropriate. Data is presented as mean ± SD. *indicates P < 0.05, ** indicates P < 0.01, *** indicates P < 0.001 and **** indicates P < 0.0001. * indicates a significant difference compared with *sparc^-/-^*. n=10-12 animals (3-6 males and 5-8 females) per group and 3 levels per animal (L4-S1).

Cortical bone parameters measured in the ROI described in **Fig.S4b**, showed a significantly lower bone volume, BV, (45%), moment of inertia (72%) (**Fig.6(m-n)**), and cross-sectional thickness (8%) (**Fig.S4e**) in *sparc^-/-^* compared with wild-type mice. Both single and combination treatments significantly increased the bone volume (48%, 67%, and 63%), the increase in moment of inertia although significant was more variable with no further improvement with the combination treatment (**Fig.6(o-p)**). Cross-sectional thickness measures revealed no significant difference between any of the groups (**Fig.S4f**). In summary, our findings suggest that oral administration of senolytics improves IVD and bone health with a more robust improvement when o-Vanillin and RG-7112 are administered as a combination treatment.

## DISCUSSION

Reducing the burden of cell senescence or selectively killing SnCs with senomorphic or senolytic drugs holds great promise for the treatment or prevention of musculoskeletal-related diseases(*59*). An elevated level of senescent disc cells is found in degenerating human IVDs(*31, 60–62*), where recent studies propose them as a hallmark and major cause of IVD degeneration(*63–66*). Clearance of SnCs has been shown to delay age-related changes in mice, and as IVD degeneration and LBP are closely linked, senolytic drugs could potentially be a promising therapy for the disease(*20, 61, 67, 68*). In this study, we show that oral treatment with RG-7112 and o-Vanillin reduced behavioural indices of LBP, reduced SASP factor release from the IVDs, and removed SnCs from the IVD and spinal cord. We also found that treatment lowered the degeneration score of the IVD, reduced the expression of pain markers in the dorsal horn of the spinal cord, and improved vertebral bone quality. Together our findings suggest that treatment with RG-7112 and o-Vanillin can reduce LBP and improve tissue health, with the most prominent effect when applied as a combination.

We previously described the two compounds, the natural compound o-Vanillin and synthetic drug RG-7112, that have senolytic effects on Sn human IVD cells. Both senolytics reduce the release of SASP factors in isolated IVD cells and intact human IVDs(*34, 35*). SnCs are heterogeneous and recent reports suggest that a combination of drugs targeting multiple pathways could improve treatment outcomes(*59, 69, 70*). o-Vanillin and RG-7112 target distinct senescence-associated networks and we found that combining them greatly enhanced the senotherapeutic effect on Sn human IVD cells(*43*).

Altogether, this prompted us to evaluate if the drugs can reduce LBP in a preclinical animal model. First, we validated that the *sparc^-/-^* mouse model, which is characterized by progressive IVD degeneration and LBP, would be a suitable model to evaluate senolytic treatment(*38*). Our results confirmed previous studies showing a higher degeneration level in knockout compared to the wild-type mice(*38, 39, 71, 72*). Using the *p16^Ink4a^*senescence marker, we found a higher density of Sn IVD cells in *sparc^-/-^* compared to age-matched wildtype animals, which was positively correlated with the degree of IVD degeneration. The observed increase in cellular senescence was further confirmed by higher mRNA expression of genes involved in cellular senescence and SASP factors in *sparc^-/-^* mice. We compared our data to the Molecular Signatures Data Base (MSigDB), Reactome, and Sen Mayo gene panels developed to identify senescence and SASP factors(*73–75*) and found all significantly enriched in *sparc^-/-^*mice. In addition, *sparc^-/-^* IVDs released elevated levels of SASP factors. In previous studies, SPARC has been shown to affect senescence and tumorigenesis, with over-expression of SPARC linked to an increase in SnCs in trabecular bone, in fibroblasts, and in enhanced tumour progression(*76–80*). In contrast, our findings demonstrate higher levels of senescence markers, SASP expression and release in *sparc^-/-^* IVDs and spinal cord, supporting this animal model for the evaluation of senotherapeutics in the context of IVD-related LBP. SnCs are heterogeneous with tissue and species-specific phenotypes, therefore, before venturing into *in vivo* experiments, we confirmed that o-Vanillin and RG-7112 could reduce SASP factor release from mouse IVDs *ex vivo.* The experiments showed a reduction in most (10/13) of the tested SASP factors. The panel of SASP factors we evaluated are described as key factors upregulated in SnCs where they contribute to paracrine senescence(*3, 13, 81, 82*).

*sparc^-/-^*mice develop age-related LBP that is well established at 7 months of age(*37–39, 44*). We chose this time point to evaluate if systemic treatment with o-Vanillin and RG-7112 reduced behavioural signs of LBP, SASP factor release, SnCs, and IVD degeneration. We further investigated if a combination treatment could improve outcomes. We treated the animals orally over 8 weeks and found a significant reduction in pain behaviour as determined by a reduction in axial discomfort and mechanical sensitivity. At the termination of the full 8 weeks of treatment, lower pain sensitivity was measured in all the behavioural tests with the combination at a higher dose being the most efficient. Together, we could show that senolytics have the potential to reduce behavioural signs of LBP. We followed in previous studies the same treatment schedule with TLR and IL8 agonists and we also found that up to 8 weeks of treatment is needed to see robust disease-modifying effects(*44, 45*). Other recent reports have demonstrated that genetic and pharmacological elimination of SnCs can improve osteoarthritis pain (OA) and delay age-related IVD degeneration(*83–85*).

LBP could in part be triggered by SASP factors released from the IVD that could potentially be reduced if the drugs reach and kill SnCs in the IVD. Our results showed that senolytic treatment reduced the release of CXCL-1, -5, 9, and 10. These chemokines have been suggested to induce ECM degradation, inflammatory response(*35, 86–88*) and senescence(*13, 89*). For example, CXCL5 promoted cellular senescence in mouse embryos resulting in implantation failure by inducing suppression of cell proliferation in trophoblast cells(*86*). In addition, microarray analysis on AF tissues from discs of aged mice treated with the senolytic drugs D + Q showed downregulation of CXCL-5 gene expression(*21*). Release of the chemokines CCL-2 and CCL-7 were also reduced by o-Vanillin and RG-7112 treatment. Elevated expression of them has previously been reported in cells from degenerating IVDs(*35, 42, 88*). Additionally, CCL2 has been reported to play a crucial role in macrophage recruitment and matrix remodelling in rheumatoid arthritis(*90*). O-Vanillin and RG-7112 treatment also reduced cytokines proposed to be secreted by SnCs, including IL-1b, TNF-α, IL-2, IL-10, and IL-6. These are all factors widely recognized as important mediators of paracrine senescence in IVD degeneration and OA(*3, 7, 13, 82, 91–101*). Moreover, VEGF-a, a SASP factor that promotes vascular infiltration, was reduced. A positive correlation and co-expression of VEGF and the senescence marker p53 have been demonstrated to promote neovascular infiltration related to IVD degeneration(*102*).

Our previous work showed an effective reduction of senescent human disc cells in response to o-Vanillin and RG-7112 treatment in ex vivo and *in vitro* systems(*34, 35*). Here we could verify that oral treatment resulted in a reduced senescence burden and improved IVD tissue health in *sparc^-/-^* mice. Previous data from other groups reported a protective effect of reducing senescence with Quercetin and ABT263 in injury-induced-IVD-degeneration in rats(*97, 103*). Similarly, selective removal of *p16^Ink4a^*-positive cells in p16-3MR transgenic mice reduced age-related IVD degeneration(*61*) and removal of *p16^Ink4a^* reduced apoptosis and attenuated SASP in mice following conditional deletion of *p16^Ink4a^*in the disc(*65*).

Similarly to our results, senolytic combination treatment of D+Q is more efficient in the treatment of idiopathic fibrosis, diabetic kidney disease and age-related diseases such as bone loss, physical dysfunction, shortening health-and lifespan(*23, 104–106*). Weekly injection of D + Q also decreases senescence burden in age-dependent IVD degeneration(*21*). The beneficial effect of reducing senescence burden and IVD degeneration in our middle-aged animals suggests that o-Vanillin, RG-7112, and their combination can slow or prevent IVD degeneration. Altogether, our data suggest that o-Vanillin and RG-7112 significantly reduce SnCs and SASP factor release when administrated systemically with a combination of the two senolytics further improving the beneficial effects.

*sparc^-/-^* mice start showing behavioural signs of LBP at 4 months of age and, would have experienced pain for 3 months before we started the treatment with o-Vanillin, and RG-7112(*38*). Cellular senescence in various cell types in the CNS has been implicated in the pathogenesis of diseases featuring pain such as osteoarthritis(*35, 84, 107, 108*). An accumulation of SnCs has been reported in neurons(*53, 55, 109*), glial cells(*52, 55, 56, 110–112*), and astrocytes(*113–117*) in humans and animals with pain. Previous studies have also linked SASP and elevated senescence in astrocytes and microglia to pain(*110, 118, 119*). Spinal microglia are suggested to contribute to the development of chronic pain while astrocytes contribute to the maintenance of chronic pain(*58, 120, 121*). We found higher expression of senescence in the dorsal horn of the spinal cord in *sparc^-/-^* mice which was reduced by both single and combination treatment. An accumulation of SnCs with age has also been reported in the spinal cord of p16 reporter mice(*122*), with an observed attenuation of the neuroinflammatory phenotype following the suppression of cells expressing *p16^Ink4a^* (*118*). *A* correlation between senescence in the spinal cord and nerve injury-associated symptoms has also been described where antihyperalgesic and anxiolytic activity was reduced after treatment with the senomorphic rosmarinic acid(*110*). We previously reported an increase in immunoreactivity for the sensory neuropeptide CGRP, the astrocyte marker GFAP, and the microglia marker CD11b in the dorsal horn of the spinal cord in *sparc^-/-^* mice when compared to wild-type mice(*44, 45, 57*). In this study, we investigated pain-related neuroplastic changes in the dorsal horn in *sparc^-/-^* mice treated with senotherapeutics. Depletion of SnCs in the spinal cord of treated mice was accompanied by decreased immunoreactivity for CGRP, GFAP, and CD11b, which could reflect decreased peripheral nociceptive signalling or a decrease in spinal neuron activity. Our results are consistent with the findings of *Du et al.,* correlating astrocytic senescence, neuroinflammation at the spinal level and signs of neuropathic pain resulting from sciatic nerve injury which was reduced by treatment with D+Q(*115*).

*sparc^-/-^* mice are known to have osteopenia, which is also a potential contributor to LBP. Recent studies report that senolytic drugs have the potential to improve age-related bone loss in some models(*24, 103, 104*). Our results showed that IVD volume and vertebral bone quality could be improved by o-Vanillin and RG-7112 treatment. Similar to our findings, ABT263 or D+Q demonstrated beneficial effects on bone remodelling in aged mice by lowering bone resorption, enhancing cortical and maintaining trabecular bone formation(*104, 123, 124*). Another study reported greater numbers of senescent osteoclasts in a mouse model of spinal hypersensitivity which was reduced by treatment with ABT263(*84*). Short-term treatment with ABT263 also showed beneficial effects on bone mass and osteoprogenitor function in aged mice. In addition to decreasing SnCs burden, ABT263 treatment decreased trabecular bone volume fraction and impaired BMSC-derived osteoblasts(*125*). In contrast, senolytic combination treatment with D+Q did not improve the vertebral bone quality of aged mice(*21*).

o-Vanillin, the main metabolite of Curcuma, has previously been evaluated as an anti-inflammatory in animal models where it showed a strong safety profile(*126, 127*). An explorative toxicology study where mice were treated orally 5 days a week for 2 weeks with 60 mg/kg o-vanillin found no sign of treatment-related toxicity(*46*). Similarly, our results showed no cytotoxicity or adverse effects caused by weekly treatment of 100 mg/kg o-Vanillin over an 8-week period. RG7112 was the first MDM2 inhibitor to enter clinical trials. Severe adverse effects were common when daily high doses needed to treat cancer were administered to patients with advanced solid tumours and hematological neoplasms (NCT00559533 and NCT00623870)(*47*). Here we used comparably much lower concentrations and found no effect in weight, mortality, or distance travelled between untreated and animals treated with RG7112 alone or in combination with o-Vanillin. Together this indicates that the doses needed to target senescence are safe, although, this must be confirmed in a full toxicological study(*46–48*). There are ongoing clinical trials with the senolytic drugs Quercetin, Curcumin, UBX0101, RG-7112, digoxin, azithromycin, and fisetin (NCT04946383, NCT04590196, NCT04129944, NCT00623870, NCT04834557, NCT00285493 and NCT04210986) evaluating senolytic drug treatment for a variety of age-related disease.

The findings presented in this study show that systemic oral treatment with RG-7112 and o-Vanillin reduces behavioural indices LBP, reduces SASP, and alleviates degenerative changes in tissues across the spine and spinal cord, with combination treatment resulting in a more robust response. This suggests that senolytics such as o-Vanillin and RG-7112 could provide novel therapies for the treatment of LBP and other painful disorders where cell senescence is implicated.

## MATERIALS AND METHODS

### Study design

This study was designed to investigate the potential therapeutic effect of two senolytics (o-Vanillin and RG-7112) for the treatment of LBP. Initial experiments were designed to determine if sparc^-/-^ mice accumulate SnCs in degenerating lower lumbar IVDs, and the upper lumbar IVDs were used to evaluate the global gene expression profiles of sparc^-/-^ and wild-type mice. A second set of IVDs was used to determine SASP factor release and to validate if o-Vanillin and RG-7112 reduced SASP release from mouse IVDs ex-vivo. We next treated sparc^-/-^ mice and evaluated pain behaviour. Grip strength and tail suspension tests were used to study axial pain, while the acetone evoked behaviour and Von Frey stimulus was used to assess radiating pain. The animals were sacrificed, and a multiplex assay was used to quantify SASP factors released from IVDs of treated and untreated animals. A second set of animals was perfused and used to investigate if the drugs depleted SnCs in the IVDs and spinal cord, and FAST staining was used to determine IVD degeneration grade after treatment. Pain signaling in the spinal cord was evaluated with markers of activated astrocytes and microglia in untreated wild-type and treated and untreated sparc^-/-^ mice. A third cohort was used to evaluate the effect of senolytic drugs on IVD volume, and bone health.

### Animals

All experiments were approved by the Animal Care Committee of McGill University following the Canadian Council of Animal Care guidelines. Age-matched male and female C57BL/6N (wild-type) and sparc^-/-^ mice were used to carry out all experiments. Sparc^-/-^ mice were developed on a C57BL/6×129SVJ background as previously described(39, 128). Mice were backcrossed and housed in a temperature-controlled room with a 12-hour light/dark cycle, 2–5 per ventilated polycarbonate, cage (Allentown), and with corncob bedding (Envigo), cotton nesting squares, ad libitum access to food (Global Soy Protein-Free Extruded Rodent Diet, Irradiated) and water. Sample size was based on previous studies to detect differences between strains and potential pharmacological effects in sparc^-/-^ mice(38, 39, 44, 45, 57, 129).

### Ex vivo evaluations

#### Bulk RNA-seq and analysis

Total RNA was extracted (QIAGEN RNAease kit) and assessed for quality (2100 Bioanalyzer, Agilent) before undergoing bulk RNA-seq analysis, generating 25 million paired-end reads (PE100) per sample on the NovaSeq 6000 platform. Raw data quality was initially evaluated using FastQC (Galaxy Version 0.74+galaxy0), and subsequent Trimmomatic (Galaxy Version 0.39+galaxy0) was employed for adapter trimming and the removal of Illumina-specific sequences. HISAT2 (Galaxy Version 2.2.1+galaxy1) was employed for read alignment to the mm10 reference genome, and gene expression quantification was performed using featureCounts (Galaxy Version 2.0.3+galaxy2). Differential gene expression analysis was conducted using Limma-voom (Galaxy Version 3.50.1+galaxy0) with Benjamini–Hochberg-adjusted p-values to control the false discovery rate. All processes on the Galaxy platform were using default parameters. A 3D Principal Component Analysis plot was generated in R to visualize sample relationships using the scatterplot3d package. Gene Set Enrichment Analysis was performed with custom gene sets from MSigDB. Multiple GSEA figures were created in R using the ggplot2, plyr, grid, and gridextra packages. Heatmap visualization of selected senescence-associated genes was created using the R package heatmap.

#### Ex vivo treatment of IVDs

To evaluate if o-Vanillin and RG-7112 can target SnCs mouse IVD cells and reduce SASP factors, isolated IVDs were treated in culture ex vivo with single or combined treatments. 9-month-old sparc^-/-^ and wild-type mice were euthanized, and four discs (L1–L5) with a cartilaginous endplate and no bony endplate were excised. Sparc^-/-^ discs were cultured for 48 h in Dulbecco’s Modified Eagle Medium (DMEM) with 1× GlutaMAX, 10 U/mL penicillin and 10 μg/mL streptomycin and one of the following senolytics: 100 μM of o-Vanillin (HO), 5 μM of RG-7112 (HR), 100 μM of o-Vanillin and 5 μM of RG-7112 (HO+HR), 100 μM of o-Vanillin and 2.5 μM of RG-7112 (HO+LR), 50 μM of o-Vanillin and 5 μM of RG-7112 (LO+HR), or 50 μM of o-Vanillin and 2.5 of μM RG-7112 (LO+LR). The disc media was changed after 48 h of treatment and replaced with DMEM with 1× GlutaMAX, 10 U/mL penicillin, and 10 μg/mL streptomycin, which was then collected and used for protein analysis. Wild-type discs were cultured for a total of 96 h, with DMEM with 1× GlutaMAX, 10 U/mL penicillin and 10 μg/mL streptomycin with a media change at the 48-hour time point. The measure of the selected SASP factors release was performed using Luminex Multiplex Assay as described below.

### In vivo evaluations

#### Randomization and blinding

Animals were randomly assigned to each treatment group.

Experimenters were blinded to genotype and treatment for all experiments and data analysis.

#### Selection of start and endpoints

The selection of treatment frequency and endpoints was based on previous studies and pilot experiments(38, 39, 44, 45, 57, 129). The start point of treatment was selected based on pain phenotype and IVD degeneration status. sparc^-/-^ mice have established back pain and IVD degeneration at 7 months of age that gets progressively getting worse(37–39, 44, 45). We aimed to determine a potential protective or regenerative effect of the senolytics either alone or as a combination treatment.

#### Animals and treatment

7-month-old_male and female mice were randomized into treatment groups. Drug doses were selected extrapolated from ex vivo studies by converting the highest optimal concentration in culture (µM) to the highest dose in vivo (mg/kg). Experimenters were blinded to the treatment groups for all experiments. sparc^-/-^ and wild-type mice underwent oral gavage every week for 2 months. Sparc^-/-^ mice received either, 100mg/kg o-Vanillin (HO), 5mg/kg RG-7112 (HR), 100mg/kg o-Vanillin and 5mg/kg RG-7112 (HO+HR), 100mg/kg o-Vanillin and 2.5mg/kg RG-7112 (HO+LR), 50mg/kg o-Vanillin and 5mg/kg RG-7112 (LO+HR), 50mg/kg o-Vanillin and 2.5mg/kg RG-7112 (LO+LR) or vehicle (0.01% Dimethyl sulfoxide (DMSO) in saline) as a control. All wild-type mice underwent oral gavage with vehicle control.

#### Pain Behaviour

Pain behaviour was performed as was previously described in our laboratory(38, 39, 44, 45, 57, 129). All mice were tested in a dedicated behavioural testing room with regular indoor lighting between 8:00 AM and 12:00 PM. Mice were habituated to the room for 1 h and to Plexiglas testing boxes on a metal grid for another hour (when applicable). Grip strength (axial discomfort), acetone-evoked behaviour and mechanical sensitivity to von Frey filaments (radicular pain) were assessed on non-treatment days bi-weekly. The tail suspension test (axial discomfort) and distance travelled in an open field (motor ability and anxiety-like behaviour) were measured every 4 weeks while mice weighing was performed every 2 weeks.

#### Grip Strength

Axial discomfort was measured with a Grip Strength Meter (Stoelting Co.) by allowing the mice to grip a bar with their forepaws and stretching them by pulling on their tail. The force at which they released was recorded in grams(130). In a session, grip strength was measured two to three times and then averaged. Mice were returned to their home cages for approximately 15 minutes between measurements.

#### Acetone-evoked Behaviour Test for Cold Sensitivity

Behavioral reaction to a cold stimulus was used for radiating leg pain. Acetone (30-50 mL) was applied to the left and right hind paws, and the total time of evoked behaviour (paw lifting, shaking, and scratching) was recorded for 30s(130).

#### Von Frey Test for Mechanical Sensitivity

von Frey filaments (Stoelting Co.) were applied to the left and right hind paw plantar surfaces until withdrawal or for a total of five seconds, whichever came first(39). Stimuli intensity ranged from 0.6 to 4.0 g, corresponding to filament numbers 3.22, 3.61, 3.84, 4.08, and 4.17. The 50% withdrawal threshold in grams was calculated using the up-down method(131).

#### Tail Suspension Assay

Mice were individually suspended by the tail underneath a platform. Adhesive tape was used on two points to attach the tail (0.5-1 cm from the base and the tip of the tail) to the platform and were videotaped for three minutes. The duration of time spent in a) immobility (not moving but stretched out), b) rearing (trying to reach the underside of the platform), c) full extension (actively reaching for the floor), and d) self-supported (holding either the base of its tail or the tape), was analyzed by a blinded observer using digital software (Labspy®, Montreal, QC) over the entire testing period.

#### Open Field

Mice were placed in a 24 cm by 24 cm Plexiglas enclosure for five minutes. Mice were video recorded from above, and the total distance travelled was analyzed using AnyMaze(*38*).

#### Luminex Multiplex Assay

SASP factor release was measured using the Luminex multiplex assay (PPX-15-MXGZF4V). *Ex-vivo* samples were prepared and collected as described above. *In-vivo* samples were obtained following treatment with senolytic compounds from *sparc^-/-^* and wild-type mice as described above. Fifteen proteins were analyzed according to the manufacturer’s instructions. Concentrations (pg/mL) (INF-γ, TNF-α, IL-1β, IL2, IL6, IL10, CCL2, CCL7, CXCL1, CXCL5, CXCL9, CXCL10, M-CSF, RANKL, VEGF-A) were measured in 40 μL media.

Median fluorescence intensity (MFI) from microspheres was acquired with a BD FACSCanto II and analyzed in FlowCytomix Pro2.2.1 software (eBioscience). The concentration of each analyte was obtained by interpolating fluorescence intensity to a seven-point dilution standard curve supplied by the manufacturer.

#### Histological analysis

##### Sample preparation

Animas were deeply anesthetized by an intraperitoneally administered mixture of ketamine 100 mg/kg, xylazine 10 mg/kg, and acepromazine 3 mg/kg, and perfused through the left cardiac ventricle with vascular rinse followed by 200 mL of 4% paraformaldehyde in 0.1 M phosphate buffer, pH 7.4, at room temperature, for 20 minutes. The T13–S1 spinal segment was collected post-perfusion and kept in fixative overnight at 4°C, and then decalcified by immersion in 4% ethylenediaminetetraacetic acid in phosphate-buffered saline at 4°C for 14 days. The samples were cryoprotected in 30% sucrose in phosphate-buffered saline for 4 days at 4°C and embedded in an optimal cutting temperature cutting medium (Tissue-Tek, Sakura Finetek, Torrance, CA, USA). Sixteen micrometre-thick cryostat (Leica CM3050S, Leica Microsystems, Inc, Concord, Ontario, Canada) sections were cut in the sagittal plane, thaw-mounted onto gelatin-coated slides and stored at −20°C.

##### FAST (fast green, Alcian blue, Safranin-O, and tartrazine) Staining

Staining was performed as described by Millecamps et al(*37–39, 44, 45, 57, 129*). This staining was adapted from the FAST method for colorimetric histologic staining described by Leung et al. for intervertebral discs. The FAST profile identifies intervertebral disc compartments and detects matrix remodeling within the disc. The procedure consists of consecutive baths in 1) Acidic Alcian Blue (Sigma #A5268), 10 minutes; 2) Safranin-O (Sigma #S2255), 2.5 minutes; 3) 50% ethanol, 1 minute; 4) Tartrazine (Sigma #HT3028), 10 seconds; and 5) Fast Green (Sigma #F7252), 5 minutes. After drying, slides were mounted with Dibutylphthalate Polystyrene Xylene (DPX) (Sigma-Aldrich, USA). IVD degeneration severity grading scale: Grade 0) Healthy IVDs display intact structure, a clear distinction between outer AF and inner NP and negatively charged proteoglycans; Grade 1) The changes in extracellular components and IVD integrity were identified as grade 0 -normal structure, but the loss of proteoglycans in inner NP; Grade 2) Internal disruption (loss of boundary) between NP and AF; Grade 3) Bulging of NP in dorsal aspect; Grade 4) Herniation.

##### Spine immunofluorescent histochemistry

Samples were prepared as described under the “histological analysis sample preparation” section. Sagittal sections (Three per animal) were incubated for 1 h at room temperature in a blocking buffer containing 0.3% Triton X-100, 1% bovine albumin, 1% normal donkey serum, and 0.1% sodium azide in PBS. Slides were then incubated with recombinant anti-CDKN2A/ *p16^Ink4a^* antibody (1:100, ab211542, Abcam) in blocking buffer overnight at 4 °C, washed three times for five minutes with PBS and incubated for 1.5 h at room temperature with secondary antibody (Donkey anti-Rabbit Cy3, catalog # 711-165-152) in blocking buffer. DAPI (1:50000 in PBS, Sigma-Aldrich) was briefly applied (5 min), and slides were washed thrice for five minutes. Coverslips were mounted using Aqua Polymount (Polysciences Inc.). Images were taken at 10× magnification using an Olympus BX51 microscope equipped with an Olympus DP71 camera (Olympus). *p16^Ink4a^* and DAPI images were merged using ImageJ, and the percentage of *p16^Ink4a^* positive cells was obtained by counting the total number of *p16^Ink4a^* positive cells over the total number of cells (DAPI counterstain). An average of approximately 5000 cells were counted per treatment group. Image analysis was performed by an experimenter blind to strain and treatment groups.

##### Spinal cord immunofluorescent histochemistry

Spinal cords were harvested following perfusion and euthanasia and fixed in 4% paraformaldehyde for 24 hours at 4°C, followed by cryoprotection using 30% sucrose solution for 24 hours at 4 °C. Samples were embedded in blocks of six spinal cords in optimum cutting temperature medium (Tissue-Tek). 16 μm cryostat (Leica CM3050S) sections were thaw-mounted on gel-coated slides and stored at 20°C until use. Three sections per animal were randomly selected, spanning the lumbar spinal cord for each antibody. Sections were incubated for 1 hour at room temperature in a blocking buffer containing 0.3% Triton X-100, 1% bovine albumin, 1% normal donkey serum and 0.1% sodium azide in phosphate-buffered saline. Slides were then incubated with either recombinant anti-CDKN2A/ *p16^Ink4a^* antibody (1:100; Abcam, catalogue #ab211542), sheep anti-calcitonin gene-related peptide (CGRP) polyclonal antibody (1:1000; Enzo Life Sciences, catalogue# BML-CA11370100, lot# 0807B74), goat anti-glial fibrillary acidic protein (GFAP) polyclonal antibody (1:1000; Sigma-Aldrich, catalogue# SAB2500462, lot #747852C2G2), or rat monoclonal anti-CD11b antibody (1:1000; BioRad, catalogue# MCA711G, lot# 0614) in blocking buffer overnight at 4°C, washed three times for five minutes in phosphate-buffered saline and incubated for 1.5 hours at room temperature with appropriate donkey-derived secondary antibodies from Jackson Immunoresearch; Donkey anti-Sheep Cy3, catalogue# 713-165-147; Donkey anti-Goat AlexaFlour 594, catalogue# 705-85-144; Donkey anti-Rat AlexaFluor 488, catalogue# 712-225-153, Donkey anti-Rabbit AlexaFluor 488, catalogue# 711-545-152, Donkey anti-Rabbit Cy3, catalogue# 711-165-152 in blocking buffer. DAPI (1:50000 in water, Sigma-Aldrich) was briefly applied, and slides were washed another three times for five minutes. Coverslips were mounted using Aqua Polymount (Polysciences Inc.). Images were taken at 10× magnification using an Olympus BX51 microscope equipped with an Olympus DP71 camera (Olympus). Using ImageJ, a region of interest was drawn around the dorsal horn, and a threshold was established to differentiate between positive immunoreactivity (ir) and background. The percent area of the region of interest at, or above, the threshold was quantified to measure *p16^Ink4a^*-ir, CGRP-ir, GFAP-ir or CD11b-ir. The average percent area immunoreactivity across the three sections from each animal was averaged and used as the value for that mouse. Image analysis was performed by an experimenter blind to strain and treatment group.

##### Micro-CT analysis

Micro-CT scans and analysis were performed as previously described(*24, 132*). High-resolution scans of fixed spines were conducted from the L4-S1 levels to evaluate the 3D structure. The scans were acquired with a spatial resolution of 15 μm using a Skyscan 1172 micro-computed tomography (micro-CT, Bruker, Kontich, Belgium). This device was equipped with a 0.5 mm aluminum filter and operated at 45 kV voltage, 220 μA current, and a 360° rotation. Image data was captured with a 0.4° increment between rotations and an average of 4 frames per image, each with an exposure time of 1.46 s. Image reconstruction was performed using the nRecon program (Version: 1.7.1.0, Bruker), and subsequent analysis was carried out utilizing CTan (Version 1.18.8.0, Bruker). CTVOX analysis software (version 3.3, Bruker) was used for the visualization of bone remodeling. The transverse cross-sectional images were then assessed for disc volume parameters and the trabecular and cortical bone morphology. Disc volume was determined by delineating a region of interest (ROI) that involved marking the boundary between the two adjacent endplates. In the case of trabecular analysis, an ROI was selected by defining the boundary between the endplate and transverse process within the vertebral body, with the reference area situated. The 3D datasets were examined to determine bone volume fraction (BV/TV), trabecular thickness (Tb. Th), trabecular number (Tb. N), and trabecular separation (Tb. Sp). For the evaluation of cortical bone characteristics, 2D assessments were conducted to ascertain cortical bone volume (BV), cross-sectional thickness (Cs.Th), and polar moment of inertia (MMI).

##### Statistical Analyses

Power analysis was set with a margin error alpha of 0.05, a confidence level of 95% and a 50% response distribution. Power analysis determined a sample size of 8-10 wild-type and 8-10 *sparc^-/-^* animals to observe significant differences (10-15 if sex differences become apparent). Data were analyzed using GraphPad Prism 9, with P ≤ 0.05 being considered statistically different. Data are presented as mean ± standard deviation. Data was assessed by two-tailed unpaired t-tests, repeated measures of one-way ANOVA, or repeated-measure 2-way ANOVA followed by Dunnet’s or Tukey’s post hoc tests as appropriate.

## Acknowledgments Funding

This work was supported by CIHR (Grant PJT-178111, Doctoral training award #476767), Arthritis Society (Grant SOG-20-0000000075, Postdoctoral fellowship TPF-19-0513, Ph.D. Salary Awards TGP-20-0000000063 and TGP-23-0000000210), The Louise and Alan Edwards Foundation’s Edwards (Ph.D. Studentship in Pain Research #2073, The Animal Behavioral Characterization (ABC) platform), Le Réseau de Recherche en Santé Buccodentaire et Osseuse (RSBO) (major infrastructure grant, Ph.D. Salary Award 20-0000000063) for their support.

## Author contributions

LH, LSS, and JOA designed the study. LH, MM^1^, HC, CG, and MM^2^ designed, performed the experiments, collected, and analyzed data. SG, KS, OWM, EC, and SL performed the experiments and prepared figures. LH, MM^1^, and HC wrote the manuscript with input from all authors. CG, JOA, MM^2^, and LSS advised on study design and supported manuscript preparation.

## Data and materials availability

All data are available in the main text or the supplementary materials.

