## Supplemental Files for "Senolytic Treatment for Low Back Pain"

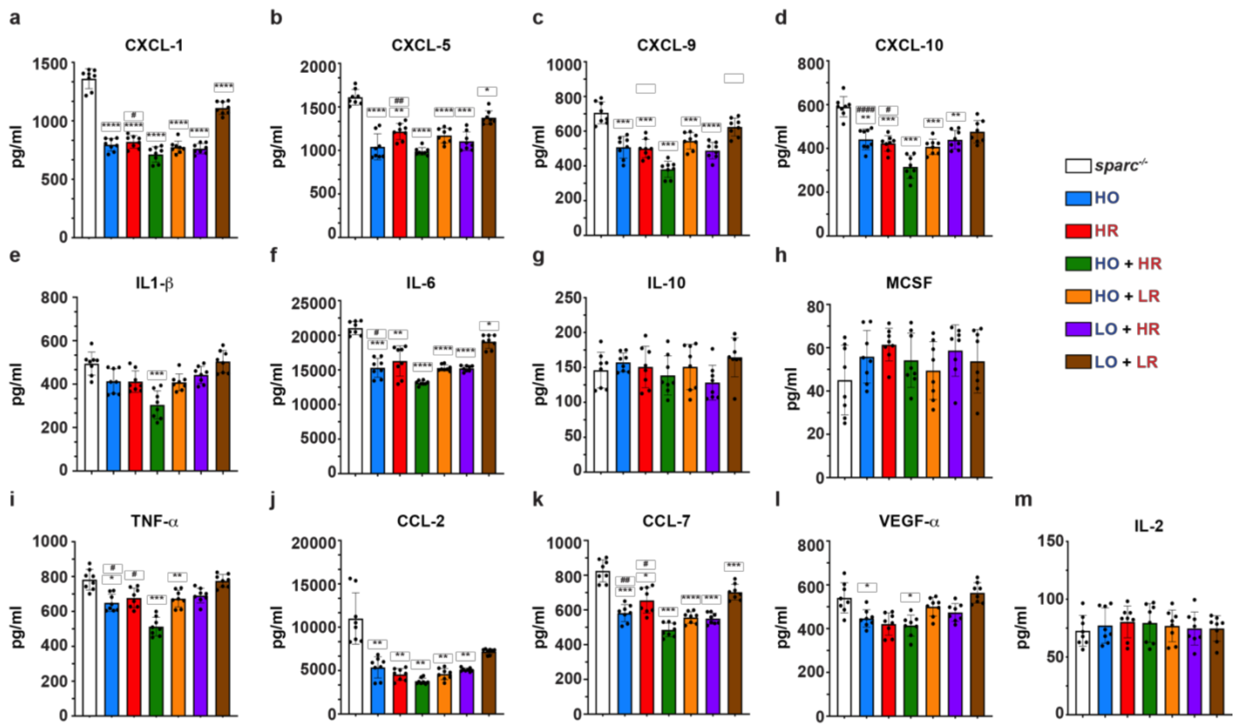


**Fig. S1 SASP factor release is reduced in *sparc$^{-/-}$* IVDs treated ex vivo.** Four IVDs were isolated (L1- L5) and treated with senolytics or vehicle and the release of 15 SASP factors was evaluated. Single treatment reluted in a significantly lower release of nine factors (CXCL-1, -5, -9, -10, IL-6, TNF-α, CCL2, CCL7, VEGF-α) (**a-d**, **f**, and **i-l**). Combination (HO+HR) showed the lowest release for the 10 factors (CXCL-1, -5, -9, -10, IL1-b , IL-6, TNF-α, CCL2, CCL7, VEGF-α) compared with single senolytics and other combinations (**a-f**, and **i-l**). No significant change was observed in the concentrations of IL-10, MCSF, and IL-2 between vehicle, single, and combination-treated IVDs (**g-h** and **m**). Means ± SD of 4 IVDs, two-way analysis of variance (ANOVA) and post hoc comparison Tukey's was used to measure significant differences between the groups. * or # indicates $P < 0.05$, ** or ## indicates $P < 0.01$, *** indicates $P < 0.001$, **** or #### indicates $P < 0.0001$. * indicates a significant difference compared with *sparc$^{-/-}$* and # indicates a significant difference compared with single drug treatment. n = 4 IVDs per animal and 8 animals (4 males and 4 females) in each group for a total of 32 IVDs per group.

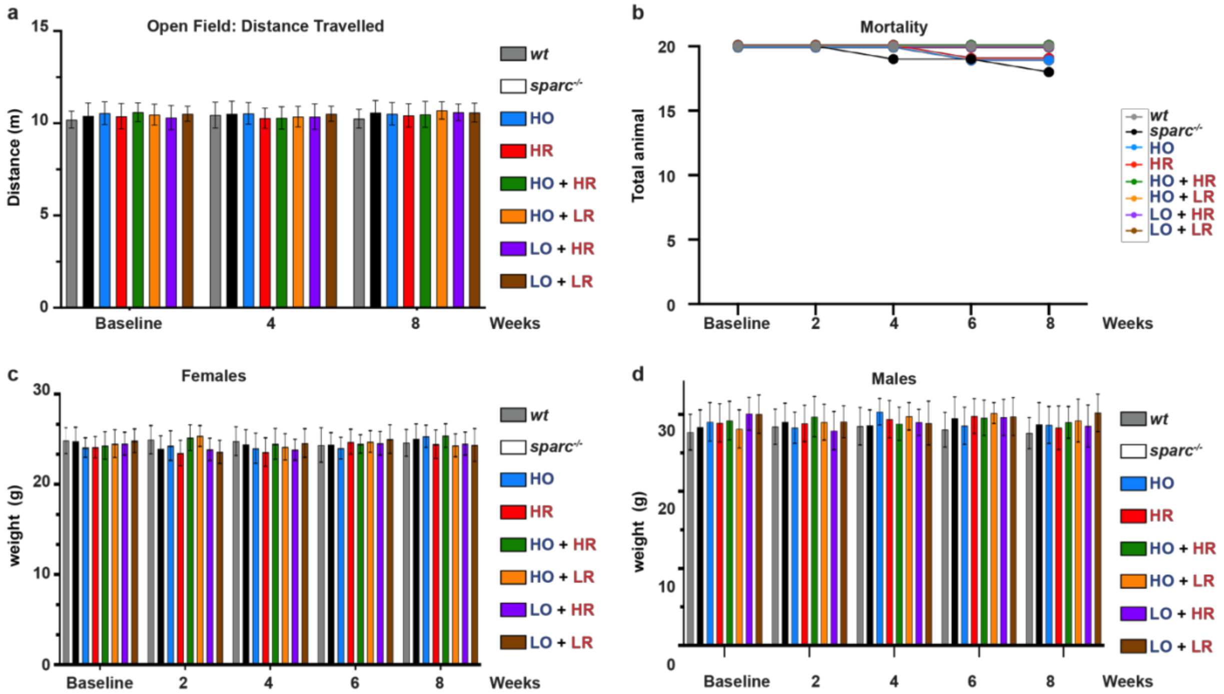


**Fig. S2 Senolytic treatment does not affect movement-evoked discomfort, mortality, and weight**. **a.** Open field (OF) test measuring the total distance covered during a 5-min in wild-type, and *sparc*$^{-/-}$ mice at baseline, 4, and 8 weeks of treatment. **b.** No significant differences were observed in mortality rates at baseline and 2, 4, 6, and 8 weeks between wildtype, *sparc*$^{-/-}$ and treated mice. **c.** Weight of female and **d.** male mice were not significantly affected by the treatment and compared with baseline at 2, 4, 6, and 8 weeks. Data is presented as mean ± SD and analyzed by one-way ANOVA followed by Tukey's post-hoc test. n = 14–20 animals (7-10 males and 9-12 females) per group, treated with senolytics or vehicle.

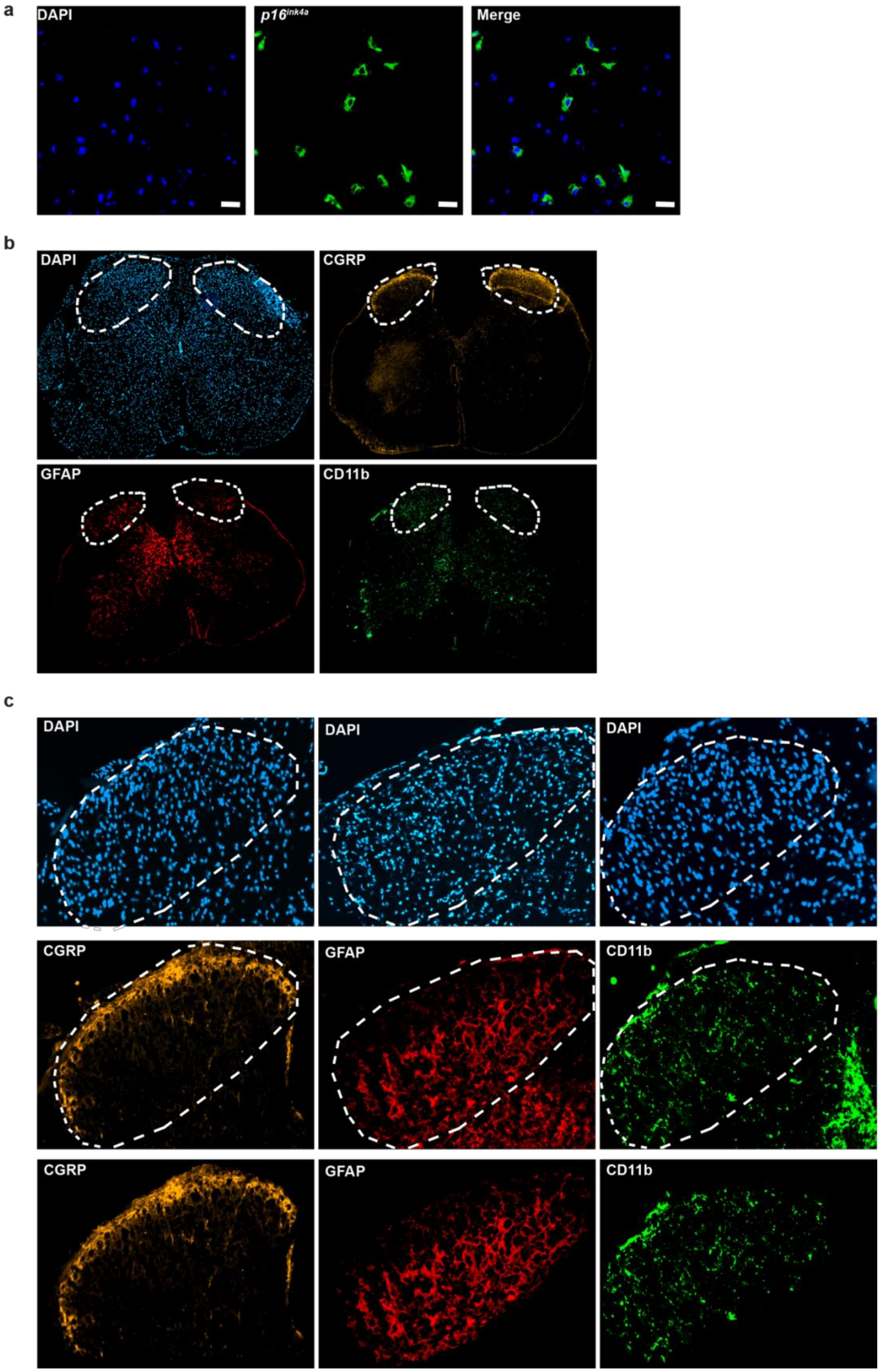
a
DAPI
p16ink4a
Merge
b
DAPI
CGRP
GFAP
CD11b
c
DAPI
DAPI
DAPI
CGRP
GFAP
CD11b
CGRP
GFAP
CD11b

**Fig. S3 Demonstration of *p16$^{Ink4a}$*-positive cells in the dorsal horn. a.** Confocal microscopy showing *p16$^{Ink4a}$* expression in the dorsal horn of *sparc$^{-/-}$* spinal cord following immunofluorescent staining. **b.** Spinal cord section images showing the ROI. **c.** Magnification of the ROIs used to identify (DAPI) and measure CGRP, GFAP, and CD11b immunoreactivity.

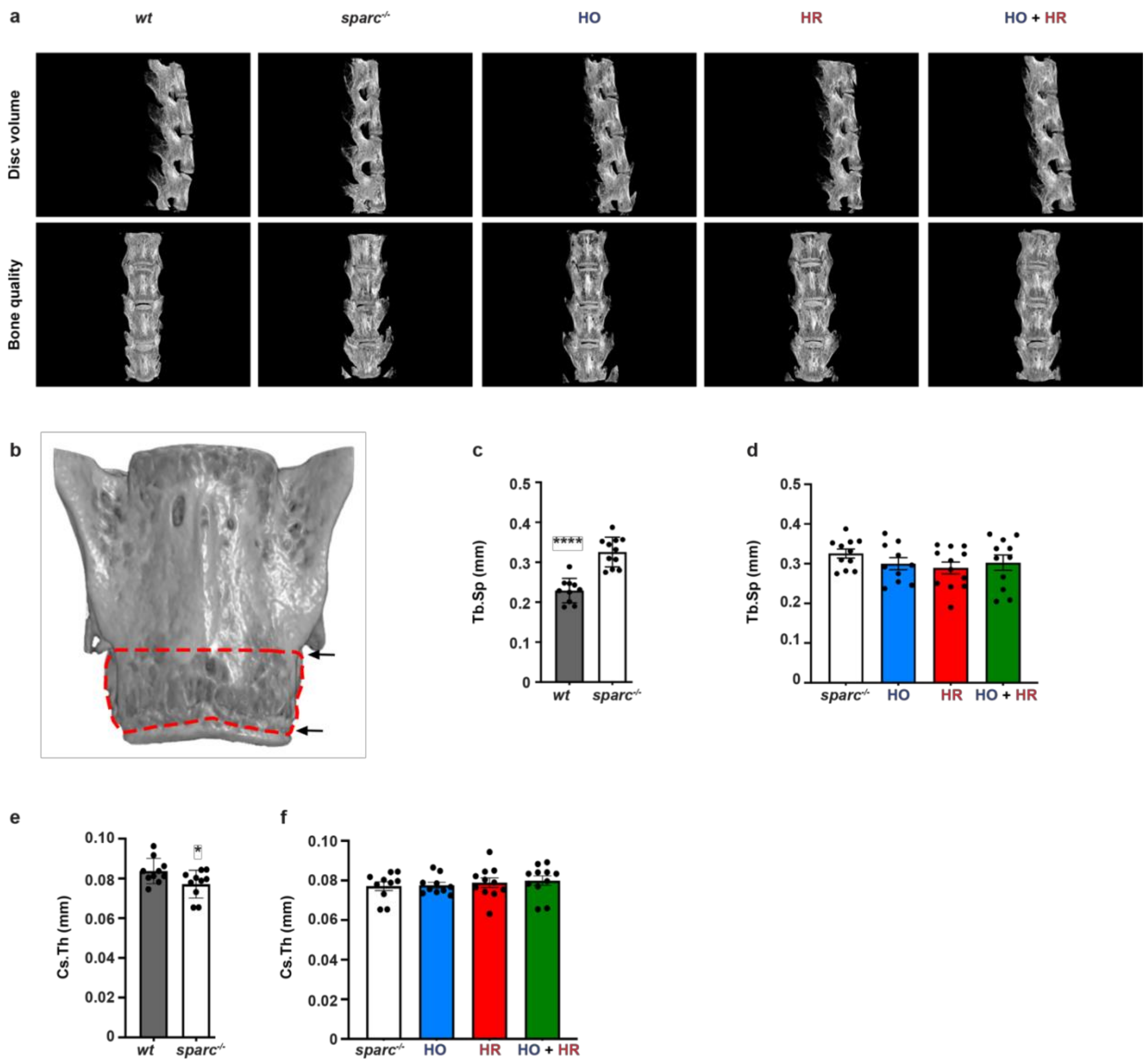


**Fig. S4 Spine reconstruction and micro-ct analysis. a.** Representative images (Videos) of wild-type and *sparc*$^{-/-}$ (vehicle or senolytics treated) illustrating the differences in disc volume and bone quality between the groups. **b**. Representative image of the ROI selected (red dashed line) to measure trabecular bone parameters. The bottom arrow indicates the endplate, and the upper arrow indicates the transverse processes that were landmarks for the ROI selection. **c**. Quantification of trabecular separation in *sparc*$^{-/-}$ and wild-type mice and **d.** Comparison of the effect of treatment in *sparc*$^{-/-}$ mice **e.** Cross-sectional thickness measure in *sparc*$^{-/-}$ and wild-type mice. **f.** Comparison of the effect of treatment in *sparc*$^{-/-}$ mice. Statistical comparisons were calculated using a Two-tailed t-test or an ordinary one-way ANOVA, with a Dunnett's post-hoc analysis as appropriate. Data is presented as mean ± SD. *indicates $P < 0.05$, and **** indicates $P < 0.0001$. n=10-12 animals (3-6 males and 5-8 females) per group and 3 IVDs/vertebra per animal. * indicates a significant difference compared with *sparc*$^{-/-}$.

**Table S1.** SASP average concentrations (pg/mL) following ex vivo treatment with senotherapeutics (Average ± STD)

| | *sparc*$^{-/-}$ | HO o-Vanillin (100 mg/kg | HR RG-7112 (5 mg/kg) | HO+HR o-Vanillin (100 mg/kg) + RG-7112 (5 mg/kg) | HO+LR o-Vanillin (100 mg/kg) + RG-7112 (2.5 mg/kg) | LO+HR o-Vanillin (50 mg/kg) + RG-7112 (5 mg/kg) | LO+LR o-Vanillin (50 mg/kg) + RG-7112 (2.5 mg/kg) |
|---|---|---|---|---|---|---|---|
| **CXCL-1** | 1360.3 ± 85 | 795.5 ± 53 | 820.5 ± 53 | 712.9 ± 65 | 774.9 ± 52 | 761.9 ± 43 | 1109.3 ± 54 |
| **CXCL-5** | 1611.2 ± 9 | 1042.3 ± 15 | 1217.5 ± 92 | 989.4 ± 44 | 1171.6 ± 85 | 1105.7 ± 11 | 1376.5 ± 78 |
| **CXCL-9** | 706.1 ± 63 | 507.6 ± 62 | 500.2 ± 53 | 380.8 ± 46 | 544 ± 47 | 488.3 ± 47 | 623.9 ± 51 |
| **CXCL-10** | 590.1 ± 46 | 441.3 ± 47 | 423.1 ± 35 | 315.6 ± 52 | 407.3 ± 34 | 439.5 ± 42 | 476.2 ± 51 |
| **IL1-β** | 494.5 ± 53 | 413.4 ± 56 | 411.7 ± 5 | 304.3 ± 64 | 408.4 ± 38 | 442.6 ± 4 | 504 ± 51 |
| **IL-2** | 72.7 ± 13 | 77.2 ± 15 | 80.4 ± 13 | 79.5 ± 16 | 76.9 ± 13 | 74.6 ± 14 | 74.6 ± 11 |
| **IL-6** | 21043 ± 95 | 15303.6 ± 13 | 16283.6 ± 22 | 13126.7 ± 32 | 15278.5 ± 43 | 15217.2 ± 4 | 19071.2 ± 96 |
| **IL-10** | 146.1 ± 25 | 157.2 ± 13 | 150.7 ± 3 | 138.5 ± 28 | 150.8 ± 32 | 128.2 ± 24 | 164.4 ± 27 |
| **TNF-α** | 782 ± 58 | 649.3 ± 5 | 676.8 ± 56 | 513 ± 53 | 673.3 ± 48 | 691.1 ± 41 | 775.4 ± 36 |
| **CCL-2** | 11004.9 ± 3 | 5372.1 ± 12 | 4555.1 ± 67 | 3778.6 ± 35 | 4602.1 ± 63 | 5091.7 ± 2 | 7199.7 ± 31 |
| **CCL-7** | 824.8 ± 63 | 581.7 ± 48 | 655.4 ± 73 | 485.1 ± 39 | 556.9 ± 33 | 549.9 ± 31 | 702.5 ± 45 |
| **MCSF** | 45.1 ± 16 | 55.8 ± 12 | 61.4 ± 7 | 54.1 ± 12 | 49.4 ± 13 | 58.7 ± 11 | 53.7 ± 14.7 |
| **VEGF-α** | 540.3 ± 68 | 446.5 ± 4 | 420.8 ± 51 | 414 ± 48 | 499.8 ± 42 | 473.8 ± 4 | 563.2 ± 46 |

**Table S2.** SASP average concentrations (pg/mL) following in vivo treatment with senotherapeutics (Average ± STD)

| | *sparc*$^{-/-}$ | HO<br>o-Vanillin (100 mg/kg | HR<br>RG-7112 (5 mg/kg) | HO+HR<br>o-Vanillin (100 mg/kg) + RG-7112 (5 mg/kg) | HO+LR<br>o-Vanillin (100 mg/kg) + RG-7112 (2.5 mg/kg) | LO+HR<br>o-Vanillin (50 mg/kg) + RG-7112 (5 mg/kg) | LO+LR<br>o-Vanillin (50 mg/kg) + RG-7112 (2.5 mg/kg) |
|---|---|---|---|---|---|---|---|
| **CXCL-1** | 1531.9 ± 13 | 974.9 ± 9 | 927.6 ± 12 | 673.9 ± 55 | 936 ± 92 | 980.2 ± 9 | 1304.4 ±34 |
| **CXCL-5** | 466.5 ± 37 | 195.6 ± 24 | 202.5 ± 33 | 124.3 ± 18 | 190.2 ± 27 | 206.6 ± 34 | 382.6 ± 22 |
| **CXCL-9** | 249.7 ± 18 | 172.2 ± 15 | 168.2 ± 16 | 144.3 ± 4 | 173.2 ± 14 | 174.8 ± 14 | 227.6 ± 16 |
| **CXCL-10** | 251.2 ± 31 | 100.7 ± 13 | 101.4 ± 13 | 73.6 ± 11 | 101.4 ± 17 | 102.2 ± 19 | 163.7 ± 8 |
| **IL1-β** | 225.3 ± 17 | 123.1 ± 13 | 121 ± 9 | 62.2 ± 6 | 123.4 ± 12 | 127.4 ± 14 | 196.5 ± 15 |
| **IL-2** | 26.8 ± 2 | 22.5 ± 2 | 26.1 ± 3 | 23.7 ± 1 | 23.8 ± 2 | 22.6 ± 1 | 23 ± 2 |
| **IL-6** | 19896.1 ± 12 | 15988.9 ± 5 | 16171.8 ± 84 | 14120.3 ± 61 | 16036 ± 73 | 15424.6 ± 34 | 17074.7 ± 56 |
| **IL-10** | 28.8 ± 3 | 25.4 ± 3 | 25.5 ± 3 | 22.8 ± 1 | 23 ± 1 | 22.5 ± 1 | 22.8 ± 2 |
| **TNF-α** | 111.7 ± 3 | 81.6 ± 7 | 85.3 ± 8 | 61.9 ± 4 | 84.3 ± 6 | 84.8 ± 8 | 95 ± 2 |
| **CCL-2** | 8576.6 ± 42 | 6720.7 ± 51 | 6427.6 ± 54 | 4732.3 ± 53 | 6529.3 ± 52 | 6584.2 ± 34 | 7235.8 ± 23 |
| **CCL-7** | 692.1 ± 72 | 463.6 ± 32 | 457.8 ± 2 | 398.7 ± 12 | 470.7 ± 23 | 466 ± 26 | 643.1 ± 54 |
| **MCSF** | 27.6 ± 2 | 25.1 ± 3 | 25.6 ± 2 | 22.4 ± 1 | 23.3 ± 1 | 23.4 ± 1 | 23.3 ± 2 |
| **VEGF-α** | 591.7 ± 45 | 455.7 ± 4 | 449.4 ± 25 | 355.1 ± 39 | 485.5 ± 47 | 491.3 ± 5 | 520.1 ± 17 |

**Table S3.** Disc and Bone parameters measurements following treatment with senotherapeutics (Average ± SD).

| Measurements | | Wild-type | *sparc*$^{-/-}$ | HO<br>o-Vanillin (100 mg/kg) | HR<br>RG-7112 (5 mg/kg) | HO+HR<br>o-Vanillin (100 mg/kg) + RG-7112 (5 mg/kg) |
|---|---|---|---|---|---|---|
| Disc Volume ($mm^3$) | | 0.73 ± 0.06 | 0.49 ± 0.16 | 0.54 ± 0.04 | 0.52 ± 0.06 | 0.62 ± 0.07 |
| Trabecular Bone | BV/TV (%) | 18.71 ± 2.76 | 8.19 ± 1.06 | 12.41 ± 3.28 | 13.81 ± 3.05 | 15.18 ± 4.68 |
| | Tb.Th (mm) | 0.064 ± 0.007 | 0.054 ± 0.006 | 0.061 ± 0.006 | 0.062 ± 0.008 | 0.067 ± 0.008 |
| | Tb.Sp (mm) | 0.23 ± 0.03 | 0.33 ± 0.04 | 0.30 ± 0.05 | 0.29 ± 0.05 | 0.30 ± 0.06 |
| | Tb.N (1/mm) | 2.96 ± 0.41 | 1.49 ± 0.10 | 2.00 ± 0.42 | 2.20 ± 0.45 | 2.21 ± 0.54 |
| Cortical Bone | BV ($mm^3$) | 0.196 ± 0.031 | 0.108 ± 0.049 | 0.160 ± 0.045 | 0.181 ± 0.028 | 0.175 ± 0.044 |
| | MMI ($mm^5$) | 0.135 ± 0.029 | 0.037 ± 0.027 | 0.087 ± 0.052 | 0.113 ± 0.037 | 0.108 ± 0.041 |
| | Cs.Th (mm) | 0.084 ± 0.006 | 0.077 ± 0.007 | 0.078 ± 0.005 | 0.079 ± 0.008 | 0.080 ± 0.008 |
